# Dynamics of SARS-CoV-2 mutations reveals regional-specificity and similar trends of N501 and High-Frequency mutation N501Y in different levels of control measures

**DOI:** 10.1101/2021.06.01.446571

**Authors:** Santiago Justo Arevalo, Daniela Zapata Sifuentes, César J. Huallpa, Gianfranco Landa Bianchi, Adriana Castillo Chávez, Romina Garavito-Salini Casas, Carmen Sofia Uribe Calampa, Guillermo Uceda-Campos, Roberto Pineda Chavarría

## Abstract

Coronavirus disease 2019 (COVID-19) is a contagious disease caused by severe acute respiratory syndrome coronavirus 2 (SARS-CoV-2). This disease has spread globally, causing more than 161.5 million cases and 3.3 million deaths to date. Surveillance and monitoring of new mutations in the virus’ genome are crucial to our understanding of the adaptation of SARS-CoV-2. Moreover, how the temporal dynamics of these mutations is influenced by control measures and non-pharmaceutical interventions (NPIs) is poorly understood. Using 1 058 020 SARS-CoV-2 from sequenced COVID-19 cases from 98 countries (totaling 714 country-month combinations), we perform a normalization by COVID-19 cases to calculate the relative frequency of SARS-CoV-2 mutations and explore their dynamics over time. We found 115 mutations estimated to be present in more than 3 % of global COVID-19 cases and determined three types of mutation dynamics: High-Frequency, Medium-Frequency, and Low-Frequency. Classification of mutations based on temporal dynamics enable us to examine viral adaptation and evaluate the effects of implemented control measures in virus evolution during the pandemic. We showed that Medium-Frequency mutations are characterized by high prevalence in specific regions and/or in constant competition with other mutations in several regions. Finally, taking N501Y mutation as representative of High-Frequency mutations, we showed that level of control measure stringency negatively correlates with the effective reproduction number of SARS-CoV-2 with High-Frequency or not-High-Frequency and both follows similar trends in different levels of stringency.

## Introduction

Coronavirus disease 2019 (COVID-19) is a contagious disease caused by severe acute respiratory syndrome coronavirus 2 (SARS-CoV-2), a single-stranded positive RNA virus that infects humans. Since the first reported cases in December 2019, the disease has spread globally causing more than 161.5 million confirmed cases and 3.3 million deaths as of May 16^th^ [1].

Since the emergence of COVID-19, significant genomic sequencing efforts have played a central role in furthering our understanding of the evolutionary dynamics of the virus. This has allowed the identification of mutations that appeared early in the pandemic (and that now seem to be fixed in the population [2,3,4,5,6]), as well as monitoring of the effectiveness of vaccines against variants coding for mutations in the spike [7,8,9,10,11,12]. Both underscore the importance of timely identification and surveillance of mutations with significant representation in the population, to efforts aimed at containing transmission of the virus

The combination of virus spread by droplets through close contact [13,14], the large number of asymptomatic cases [15,16], the absence of effective pharmaceutical treatments at the beginning of the pandemic and the delays in production and distribution of vaccines [17], leave non-pharmaceutical interventions as the most effective measures to contain the spread of COVID-19 for a large fraction of the world’s population.

Different studies have evaluated the relationship between non-pharmaceutical interventions (NPIs) and the decrease in the number of cases [18,19], the reproductive number [18,20,21], the case fatality rate [22], the contagion rate [23], and the number of SARS-CoV-2 importations [24,25]. By contrast, the effect of NPIs on specific mutations has been less well-studied. Pachetti et al. (2020) [22] analyzed how lockdown policies might have influenced the dynamics of some SARS-CoV-2 mutations; however, results are primarily qualitative and little quantitative description of the reported effect is provided. Muller et al. (2021) [26] use phylogenetic methods to estimate the importance of SARS-CoV-2 introductions on increasing the relative frequency of the D614G mutation, implicitly showing that international movement can affect the relative frequency of mutations.

Here, using 1 058 020 genomes from sequenced COVID-19 cases, we analyze the temporal dynamics of SARS-CoV-2 mutations estimated to be present in more than 3 % of global COVID-19 cases. We then investigate whether mutations are region-specific and if there is a correlation between level of lockdown policies and the effective reproduction number of specific mutations.

## Results and discussions

### 115 mutations overpass presence on 3 % of global COVID-19 cases and most of them are non-synonymous

We performed a by case normalization of the frequencies of the mutations from 1 058 020 genomes all around the world. The relative frequency of cases where a mutation is present was named Normalized Relative Frequency of a genomic position: NRFp. The NRFp of each mutation was calculated from genomes and the number of cases of 714 country-month combinations, including 98 countries from January 2020 to April 2021.

This normalization allowed us to identify mutations that have not been reported in other global studies, such as that of Castonguay et al. 2021 [27]. This is because in many countries the number of sequenced genomes is low and certain mutations could go unnoticed. Thus, we identified 115 mutations with NRFp > 0.03 (Fig. S1); this means that those mutations are estimated to be present in more than 3 % of the COVID-19 cases globally. Considering that the sum of the reported cases from the 714 country-month combinations analyzed was 120 008 410 cases, an NRFp of more than 0.03 means that those mutations were present in more than 3 600 252 global COVID-19 cases.

Table S1 summarizes the features of these 115 mutations. Based on those 115 mutations, we calculated a dN/dS ratio of 4.1 that could imply positive selection occurring in the SARS-CoV-2 genome. Additionally, S and N proteins did not show synonymous mutations and presents ∼74 % of the total non-synonymous mutations suggesting that positive selection is predominantly in those two ORFs.

### Mutations show three types of temporal dynamics

The dynamics of the 115 mutations were analyzed through calculating the NRFp in each month from January 2020 to April 2021 (Fig. 1). We assigned type of temporal dynamics to the mutations according to the NRFp in different months and the change of NRFp between months. Thus, three types of temporal dynamics were observed: i) High-Frequency mutations (HF) that never show negative NRFp changes greater than 1 %, and increased rapidly in NRFp since their appearance (Fig. 1a), ii) Medium-Frequency mutations (MF) that alternates between negative and positive NRFp changes and presents at least one month with NRFp greater than 15 % (Fig. 1b), and iii) Low-Frequency mutations (LF) that also have an alternation between negative and positive NRFp changes but at a NRFp ever below 15 % (Fig. 1c).

**Figure 1.**
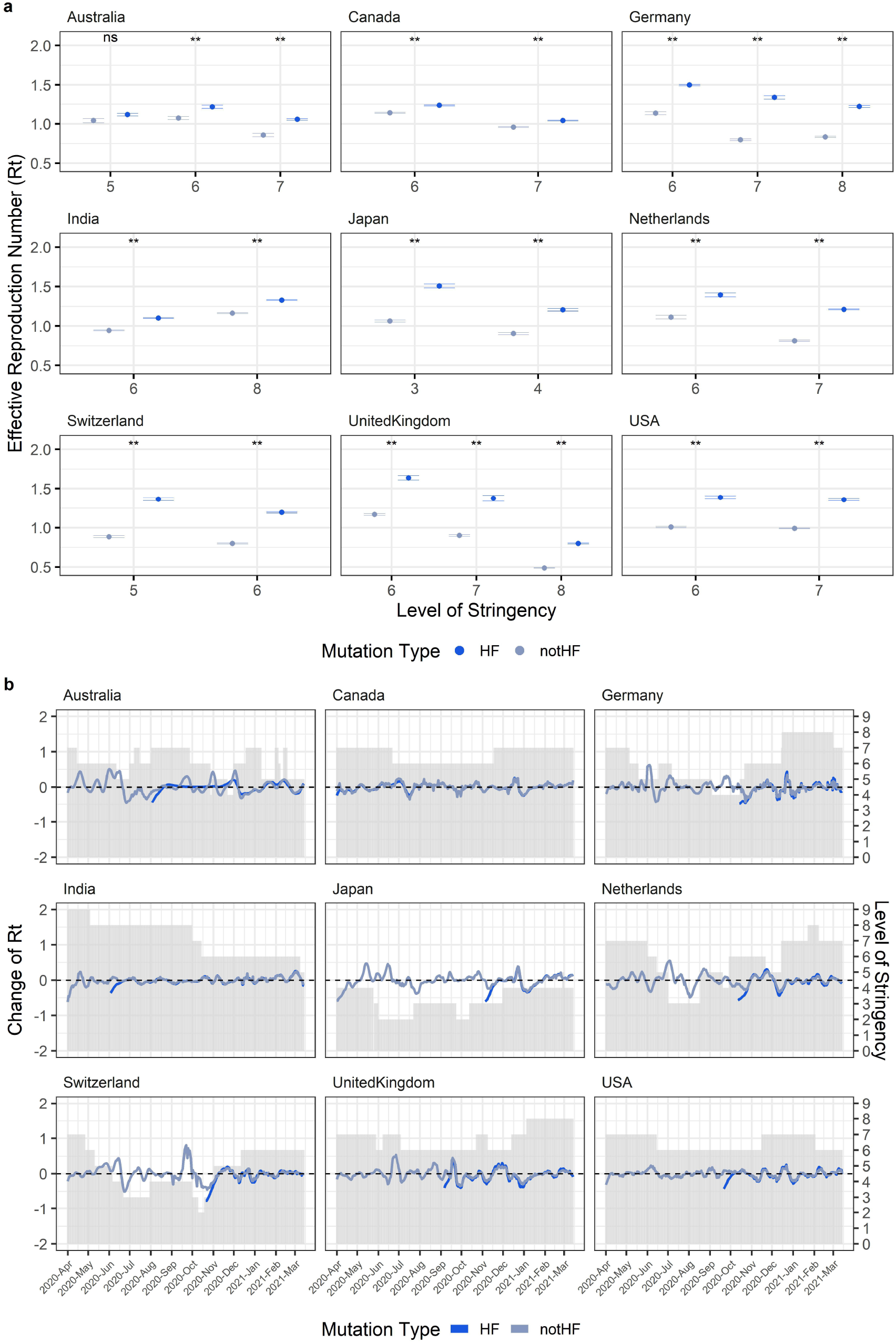
Three different temporal dynamics of SARS-CoV-2 mutations. Normalized by cases Relative Frequency (NRFp) of the mutations by month. **(a)** High-Frequency mutations (HF) never show negative NRFp changes greater than 1 %, and increased rapidly since their appearance. **(b)** Medium-Frequency mutations (MF) alternates between negative and positive NRFp changes and presents at least one month with NRFp greater than 15 % **(c)** Low-Frequency mutations (LF) that also have an alternation between negative and positive NRFp changes but at a NRFp ever below 15 %. Error bars represent inter-region variation as weighted variance.

HF mutations are characterized by a rapid increase in global frequency following their appearance (Fig. 1a). This could be due to positive selection without competition and/or by other effects related to population dynamics such as control measures implemented by countries aimed at controlling transmission. Mutations in this category appeared in two well-defined stages of the pandemic. The first group is composed of four mutations that now appear to be globally fixed. They emerged at the beginning of the pandemic in January 2020, reaching more than 0.75 NRFp in April 2020 (Fig. 1a, Group 1). The second group rapidly increased in frequency in December 2020, and have continued to increase since then (Fig. 1a, Group 2).

Some HF mutations identified here have been widely reported [28] due to their presence in variants of concern. The first and second groups contained Spike mutations well known due to their possible implications in transmissibility, (e.g. D614G in the first group [29,30] (Fig S2b)), and vaccine efficacy (e.g. Δ69-70, N501Y, and E484K, all present in the second group [31,32] (Fig. S2b and S2e)). In the future, analysis of the dynamics of other mutations in this way could help facilitate rapid identification of other mutations of concern.

By contrast, some of the MF and LF mutations that we observed have not been less previously reported to a significant degree, with descriptions either limited to specific countries or regions [33,34,35], or not reported at all, (e.g. K997Q on nsp3 and S202C on N protein). However, those mutations are present in several months throughout the pandemic and we did not observe evidence of the extinction of any of these mutations (relative frequency of 0 or near to 0 in two or more consecutive months) (Fig. 1b and 1c)

One possibility for the existence of MF and LF mutations is that some benefits may be conferred to SARS-CoV-2 but competition with other variants prevents rapid increases in their frequency increase across the population. Such dynamics have been observed in evolution experiments for other organisms [36,37]. Furthermore, the coexistence of different lineages of the same organism in the context of frequency-dependent interactions has been reported in yeast [38] and bacteria [39,40], and have highlighted that this can be beneficial for the organism. In the case of virus, epitope diversity and host-specific adaptation can be beneficial for the viral population [41].

### Some of the MF mutations are region-specific while other have medium frequencies in various regions

In our previous work [42], we observed that the mutation T85I in nsp2 has a higher frequency in North America than in other continents. Here, we show that this MF mutation maintains this tendency, persisting since its appearance at a global NRFp of ∼0.2 (Fig. S3a). Interestingly, and in contrast to HF mutations (that are typically similarly frequent across several analyzed regions, with an exception being a group of recent mutations that are more frequent in South America (Fig. S4 and S5)), most of the MF mutations (18 of 29) analyzed here are most frequent in a specific region (Fig. 2).

**Figure 2.**
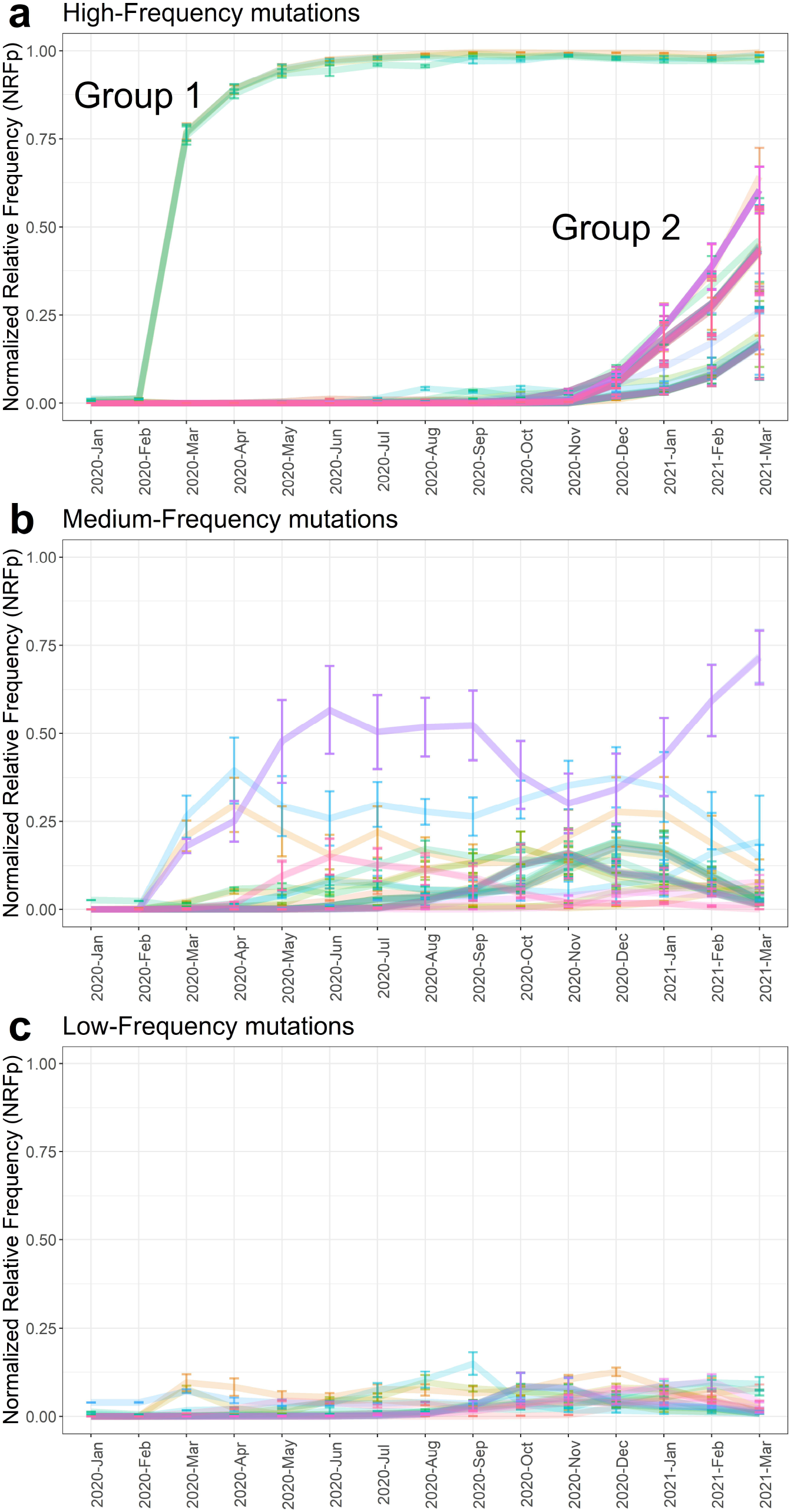
MF mutations are region-specific or have mid-frequencies in several regions. **(left-column)** Normalized by cases Relative Frequency (NRFp) of the mutations by month separated by regions (green = Africa, red = Asia, blue = Europe, grey = North America, purple = Oceania, yellow= South America). **(middle-column)** Total NRFp by region of the analyzed Medium-Frequency mutations. Numbers in each bar represents the estimated total number of cases of the particular mutation in that region. **(right-column)** Chi-square p-value and Pearson residuals analysis of Medium-Frequency mutations. Upper line corresponds to the mutant state and the bottom line to the not-mutant state. Grey and red boxes mean negative or positive association with the state, respectively. Intensity of the colors means higher residuals that means greater contribution. **(a-e)** Region-specific Medium-Frequency mutations and **(f-j)** not-region-specific medium-frequency mutations.

To explore whether MF mutations showed a region-specific pattern, we analyzed the dynamics of ten subtypes of MF dynamics in six different regions (Africa, Asia, Europe, North America, Oceania, and South America) (Fig. S6). Our results show that five subtypes had a NRFp greater than 0.3 for at least three consecutive months in only one region (Fig. 2a-e, left column). Relatedly, mutations belonging to these subtypes had a higher relative number of cumulative cases (NRFp) in a specific region, compared to other regions (Fig. 2a-e, middle column).

Then, we examined whether the proportions of estimated COVID-19 cases caused by MF mutations were different between regions. Chi-square p-values showed that in all the MF subtypes at least one region have different proportions (Fig.2, right column). Pearson residuals analysis showed which of the regions have larger or smaller mutant proportion than expected (meaning positive or negative association, red and grey squares, respectively) and which region has a greater degree of association (color intensity). The five subtypes that showed region-specific patterns also showed that just one region is positively associated and that it has the highest degree of association to that specific mutation (Fig. 2a-e, right column). By contrast, other five subtypes showed positive association to more than one region with a variety of degrees of association (Fig. 2f-j, right column)

We further analyze the five subtypes that showed region-specific pattern (Fig. 2a-e). Country analysis of the relative frequencies (Fig. S7 and S8) and the cumulative number of cases (Fig. S9 and S10) showed that those mutations are found in more than one country of the region. Some of them follows a similar pattern of frequency changes in two or more countries within the region (S7b, S7d and S8), whereas others have a particular pattern of frequency change in one particular country (S7a and S7c). Analysis of the cumulative number of cases by country showed that, although several countries present COVID-19 cases of the particular mutations, in most cases few countries contributes to most cases (S9a, Brazil; S9b, Argentina, Brazil, Chile; S9c, USA; S9d, Canada, Mexico, USA; S10, Italy, Spain, UK)

A decline of the frequency can be seen for some MF mutations in the last months (Fig. 2b, 2c, 2d), this can be explained because new mutations leave out of competition those mutations, or due to a delay between the collection date and the submission date of genomic samples. Using genomic data from August 10^th^ 2021, we re-analyzed three mutations that clearly showed this decline (Fig. 2b (I33T), 2c (A222V), and 2d (P67S)). We found very similar patterns in the countries analyzed (Fig. S11-S12), therefore, leave out of competition by other mutations is a more plausible scenario.

LF mutations followed similar patterns to those observed for MF mutations (Fig. S13 and S14). Thus, the MF and LF dynamics seems to be due to: i) high prevalence of mutations in specific regions, ii) globally dispersed beneficial mutations in constant competition with other variants, or iii) a combination of these two effects.

### SARS-CoV-2 carrying HF_N501Y_ mutation follows similar trends than SARS-CoV-2 without HF_N501Y_ in different levels of control measures

The rapid increase in global frequency of HF mutations and the observation that those mutations appeared at two very defined stages of the pandemic (Fig. 1a) lead us to hypothesize that, at least part of this abrupt increase is due to the fact that limited or minimal levels of control measures and NPIs may permit that HF mutations to spread even faster than not-HF mutations that when stronger control measures and stronger NPIs are present. An alternative hypothesis could be that strict control measures give a large competitive advantage to more transmissible variants (HF mutations), enabling them to persist and continuing to transmit, whilst their less transmissible counterparts (not-HF mutations) die out.

To test these hypotheses, we analyzed whether different degree of control measures could affect differently to SARS-CoV-2 genomes bearing the HF mutation N501Y (HF_N501Y_) or not bearing the HF mutation N501Y (not-HF_N501Y_) in nine countries that have more than 15 sequenced genomes per week during March 2020 to April 2021. We selected this mutation because it is present in three variants of concern (B.1.1.7, B.1.35, and P.1) [43] and is a good example of the behavior of HF mutations (Fig.1a and S2a). Additionally, and in contrast to HF mutations belonging to group 2, mutations in the first group of HF mutations (Fig.1A, group 1) may have been aided by founder effects in the early stages of the pandemic. For this reason, we did not analyze them in this part of our study.

First, we estimated the effective reproduction number (Rt) of HF_N501Y_ or not-HF_N501Y_ (Fig. 3a) and measure the correlation with the level of stringency (Fig. 3b). The level of stringency is a measure of the level of control policies based on nine response indicators including school closing, workplace closing, cancel public events, restrictions on gathering size, close public transport, stay-at-home requirements, restrictions on internal movement, restriction on international travel and public information campaigns [44].

**Figure 3.**
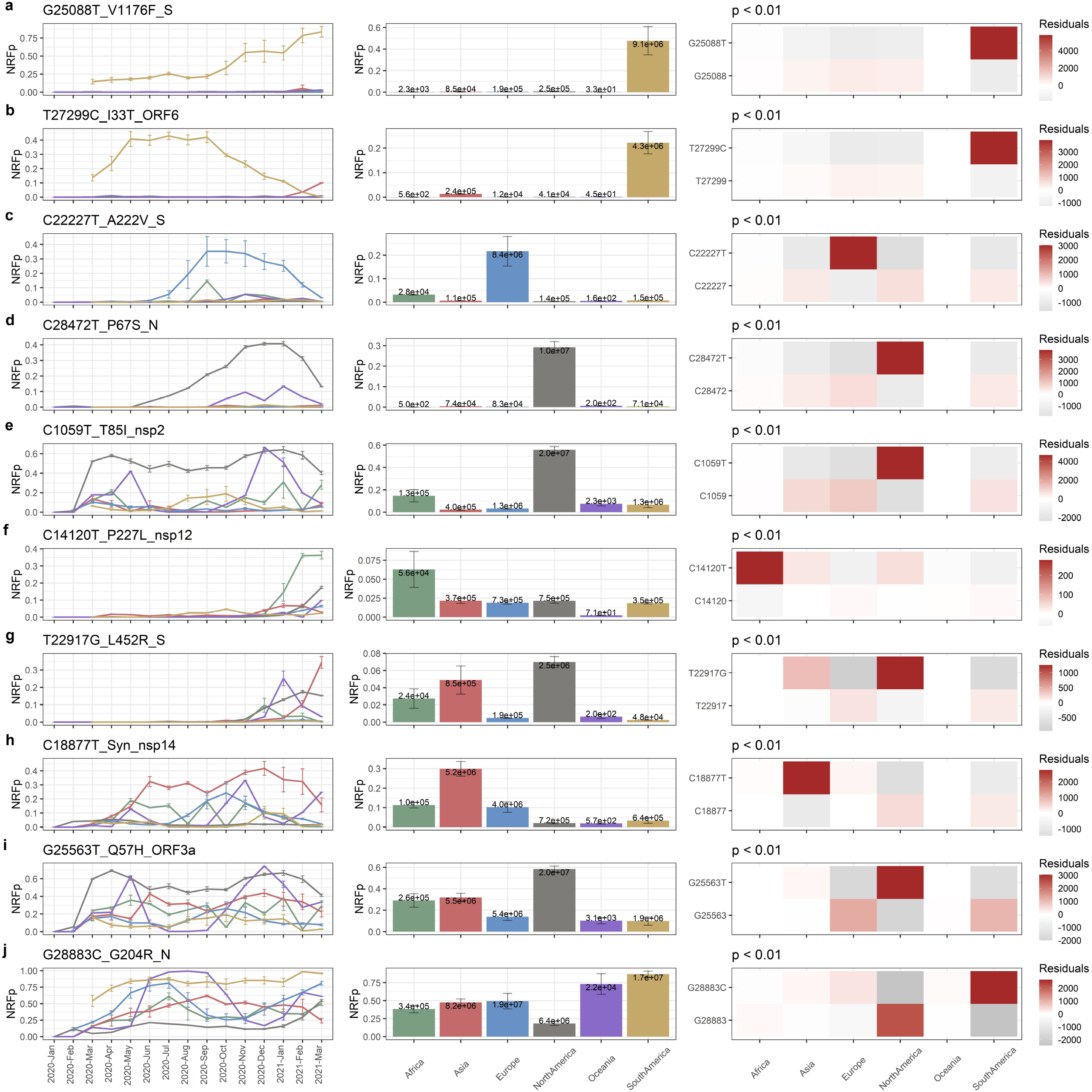
Effective reproduction number (Rt) of HF_N501Y_ and not-HFN_501Y_ are correlated with level of stringency. **(a)** Each panel shows the estimated effective reproduction number of SARS-CoV-2 bearing (blue) or not (grey) the HF mutation N501Y (HF or notHF, respectively) in different countries. Grey bars are showing the level of stringency. Shades show a 97.5 % confidence interval in the estimation of Rt. **(b)** Correlation of Rt after 14 days of the implementation of the level of stringency with the level of stringency. Each panel shows the independent analysis of different countries. Spearman correlation values (R), R-square of the linear regression model (R^2^), and p-value of the correlation is showed in the left-up of each panel in this order. Colors represent the same as in (a).

We found significant negative correlation between the Rt after 14 days that the level of stringency was implemented and the level of stringency in eight of the nine countries analyzed (Fig. 3b). In all these eight countries linear regression model explained at least 23 % of the variance in the Rt of HF_N501Y_ (Fig. 3b), and the effect size measured by the R-value of spearman correlation showed in the worst case a value of 0.48, with all the others R-value between 0.5 and 0.81 (Fig. 3b). In the case of India, the Rt of HF_N501Y_ showed a positive correlation with level of stringency. It is known that efforts in molecular testing in India have changed during the pandemic [45] Time-varying differences in the intensity and capacity of molecular testing can produce significant biases in the estimation of Rt. Overall however, our results show a significant negative correlation between degree of control measure stringency and Rt in eight of the nine countries analyzed.

We also found that, independently of the level of stringency imposed, the Rt of HF_N501Y_ was significantly higher than not-HF_N501Y_, potentially explaining why HF_N501Y_ increase its frequency faster than not-HF_N501Y_ since its appearance in the nine countries considered here (Fig. 4a). Interestingly, when we analyzed the Rt of SARS-CoV-2 genomes bearing an MF mutation (MF_R203K_) and compare it with the Rt of genomes without the MF mutation (not-MF_R203K_) we observed that in some stages of the pandemic the Rt of MF_R203K_ is higher than not-MF_R203K_ but in other cases the opposite was observed (Fig. S15). This explains why this mutation did not increases its frequency steadily and can be an evidence of constant competition between MF_R203K_ and not-MF_R203K_.

**Figure 4.**
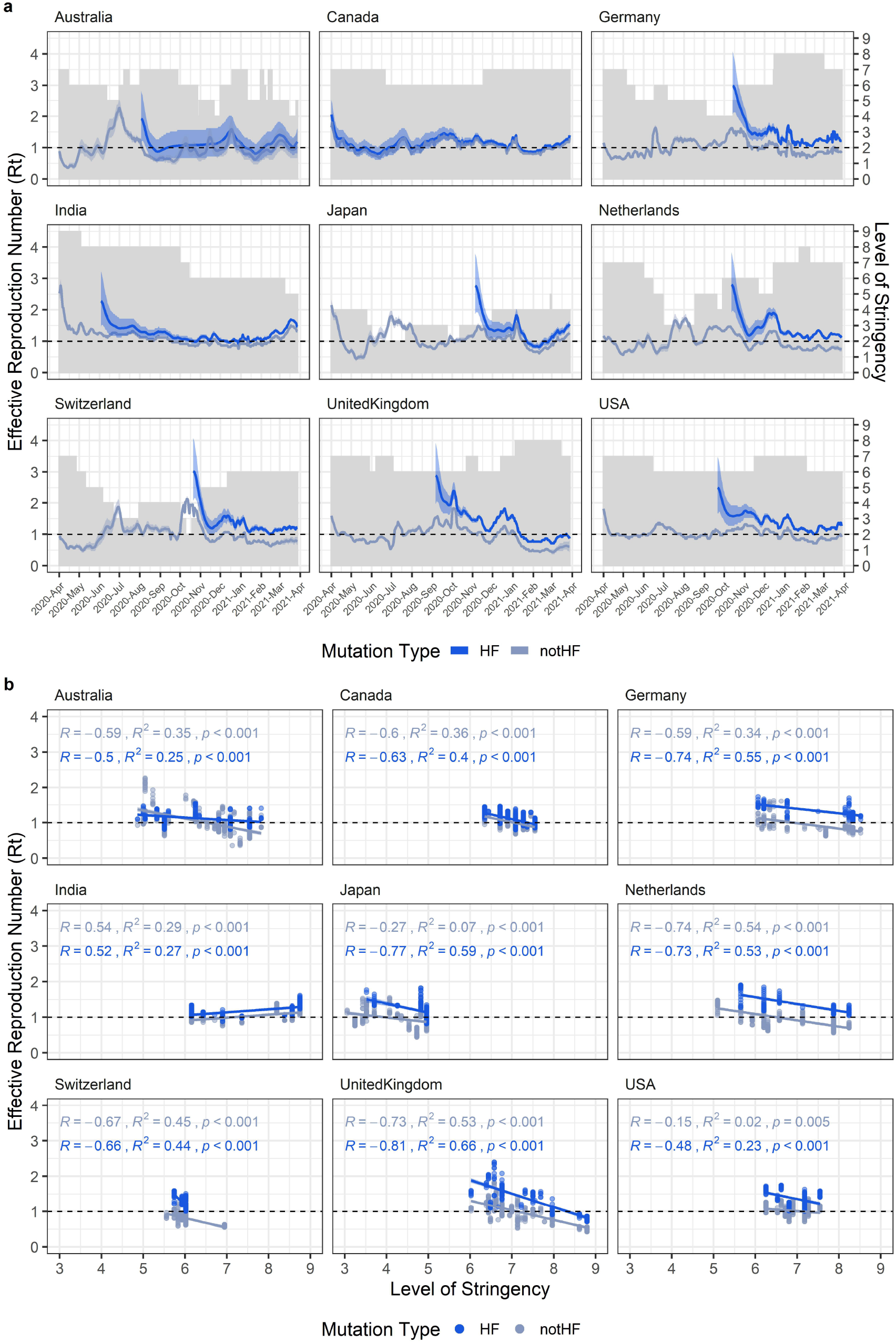
Effective reproduction number (Rt) of HF_N501Y_ is higher than not-HF_N501Y_ but similarly affected by the level of stringency. **(a)** Statistical comparison between the bootstrap distribution of the Rt of SARS-CoV-2 bearing (blue) or not (grey) the HF mutation N501Y (HF or notHF, respectively) in different levels of stringency. Points represent the mean and the lines represent the 25 and 75 percentiles of the bootstrap distribution. ** means p-value lesser than 0.05 and ns means p-value higher than 0.05. **(b)** Plot of the change of Rt in the time. Change of Rt was calculated as the Rt fourteen days after the day of interest subtracted to the Rt mean between the day of interest and thirteen days after that day. Grey bars are showing the level of stringency. Colors represent the same as in (a).

Finally, to explore whether different stringency levels differentially affect the dynamics and transmission of HF_N501Y_ and not-HF_N501Y_, we calculated the change in Rt during several months where different levels of stringency were implemented (Fig. 4b). The patterns of the change of Rt between HF_N501Y_ and not-HF_N501Y_ were almost identical (Fig. 4b) and R values of spearman correlation of HF_N501Y_ and not-HF_N501Y_ with Rt were similar in most cases (Fig. 3b), indicating that both could be similarly affected by the changes in stringency levels.

Taken together, although HF_N501Y_ presented higher Rt in lower levels of stringency indicating that HF_N501Y_ spread was likely helped by mild lockdown policies in some stages of the pandemic, this effect was also observed in not-HF_N501Y_. In conclusion, the results of this section showed control measures and their associated stringency probably affecting HF_N501Y_ and not-HF_N501Y_ in a similar fashion; thus, our two initial hypotheses are not supported by these results. Instead, the rapid increase of frequency of HF_N501Y_ is justified primarily by its generally increased transmissibility (i.e. a higher Rt which is always greater than the Rt of not-HF_N501Y_), rather than the implementation of specific control measures.

### Limitations of the study

Stringency level is calculated from set of policies applied in each country that do not necessarily operate or function the same in different countries due to, for instance, variations in sociocultural and economic factors. Thus, comparisons at country level have variation that limit the reliability and interpretability of the results presented here, especially when compared with other countries. Moreover, different combinations of policies can generate the same level of stringency - the fact that several policies were applied together to generate a stringency index precludes efforts to evaluate the effect of a specific policy on the effective reproduction number of SARS-CoV-2.

After control measures are implemented (reflected as an increase to the stringency index) Rt changes from a higher value to a lower value. This process generates a time-window of intermediate Rt before the Rt reach a plateau that indicates how much the policy lowered the Rt. These intermediate values of Rt introduce a bias in the correlation between Rt and the level of stringency. Furthermore, if a country changes the stringency level in time-windows less than those necessary for the Rt to stabilize, the estimations of correlation get more complicated.

Our correlation analysis showed that in seven of the nine countries analyzed lower levels of stringency are correlated with higher Rt values. This could be an evidence of a possible effect of lockdown policies in the Rt. However, causal inference model is known to be a more accurate approach to test causality.

Although the methodology of normalization by cases alleviates the differences in the number of genomes sequenced by country, confidence in the calculation of relative frequencies of mutations is still low in regions with a low number of genomes sequenced. For example, a mutation with 0.5 relative frequency that comes from a sample of 15 genomes will have a confidence interval between 0.25 and 0.75; on the other hand, a sample of 150 genomes will generate a confidence interval between 0.58 and 0.42. Also, the number of cases is still subjected to bias due to for instance, the difference in the number of tests that each country performs, as occurs in India.

## Conclusions

Normalization by cases of the frequency of mutations is an important tool for global analyses in a pandemic where not all the countries possess the same capacity to sequence SARS-CoV-2 genomes. This process partially mitigates differences in available genomes, but does not eliminate this problem. Worldwide efforts to help countries with fewer sequencing resources would improve our understanding of the adaptation and evolution process of SARS-CoV-2.

Three types of dynamics of mutations are described here and named “High-Frequency” (HF), “Medium-Frequency” (MF), and “Low-Frequency” (LF). The three types are represented in all the months analyzed, and found in non-structural and structural proteins, and synonymous and non-synonymous mutations. Differences in the dynamics could be due to different forces acting on each of these types of mutations and the implications of all of them need to be studied to better understand the adaptation process of SARS-CoV-2.

Medium and Low-Frequency mutations maintain roughly constants global frequency due to their higher prevalence on specific regions and/or because they are in constant competition with other mutations in several regions. We showed some mutations with a high degree of region-specificity and others that presented mid-frequencies in several regions. Higher prevalence in specific regions may be due to specific-host characteristics. Constant competition in several regions may be due to the fact that they are beneficial mutation in the presence of other mutations with a similar degree of benefit. Some mutation can be leave out of competition when others beneficial mutations appear. Our analysis, also shows evidence that some MF mutations have a reduced relative frequency after several months of high frequencies in a specific region.

In this pandemic, human behavior has strongly affected the adaptive process of the SARS-CoV-2 through continuous implementations and changes to implemented control measures. Our analysis presents evidence that the High-Frequency mutation N501Y is more transmissible (showed for its greater effective reproduction number) than not-N501Y, but also that control measures do not significantly favor the growth of any one in particular. Instead, we observe that policies have a similar impact on both.

## Methods

### Normalized by cases relative frequency of mutations on the SARS-CoV-2 genome

To perform mutation frequency analysis considering the number of cases in each country we followed similar steps as described in Justo et al. 2021 [42], with some modifications: we first downloaded 1 221 746 genomes from the GISAID database (as of April 24th, 2021). Sequences with less than 29 000 nt were removed and the resulting sequences were aligned against the reference SARS-CoV-2 genome (EPI_ISL_402125) from nt 203 to nt 29674 using ViralMSA.py [46,47]. From this alignment, we removed sequences with more than 290 Ns, more than 0.05 % unique mutations, and/or more than 2 % gaps. After those filters, we had 1 058 020 genomes. Subalignments were generated by grouping sequences by country and month. Subalignments with less than 15 sequences were not considered in the analysis. Nucleotide relative frequencies of each genomic position on each of 714 subalignments each corresponding to a different country-month combination (including 98 countries) were calculated. Normalized relative frequencies (NRFp) were calculated as the weighted mean of the relative frequencies in each subalignment with the number of cases as the weight. The number of cases for each month and country was obtained from the European Centre for Disease Prevention and Control (https://www.ecdc.europa.eu/en/publications-data/download-todays-data-geographic-distribution-covid-19-cases-worldwide). The NRFp is an estimation of the percentage of global COVID-19 cases where a particular mutation is present. The same procedure was done to obtain the NRFp of the mutations by months or by regions. Data manipulation was done using R and python scripts.

### Analysis of region-specific mutations

The frequencies by country-months of each mutation were obtained from the previous calculation. Then, the Normalized relative frequencies (NRFp) by region (Africa, Asia, Europe, North America, Oceania, South America) were calculated as the weighted mean of the relative frequencies of each country-month belonging to a specific region using the number of cases as the weight. Number of cases with a particular mutation in each country was estimated by multiplying the relative frequency of the mutation with the number of cases in a specific country-month. Then, we added the cases belonging to a specific region and chi-square analyses were done using R software [48].

### Estimation of effective reproduction number of SARS-CoV-2 mutations

We select nine countries (Australia, Canada, Germany, India, Japan, Netherlands, Switzerland, United Kingdom, USA) with at least 15 sequenced genomes by week from March 2020 to March 2021. Raw number of cases by days were obtained from [49] and used to estimate the number of cases by day for a specific mutation. In the case of MF mutation R203K, R203, and N501, we multiply the relative frequencies of the genomes with the state of interest (R203K, R203 or N501) in a determined week by the number of cases in the day. For instance, if one week presented 30 % of genomes with the mutation R203K, and the number of cases on Monday of that week was 100. Thus, the estimated number of cases with this mutation in that day was 30. In the case of the HF mutation N501Y we first calculated the relative frequencies of that mutation in each week and then adjusted the relative frequencies to a logistic regression model using R software [52]. The number of cases estimated for the MF and HF mutations by day were used to estimate the effective reproduction number using EpiFilter [50].

### Correlation analysis between stringency levels and effective reproduction number

The stringency index by country by day was obtained from [49]. Analysis of Spearman correlations and linear regression models of the effective reproduction number 14 days after the level of stringency was implemented with stringency index in each country by each state (mutant or not mutant) was done using R [48] and the packages ggplot2 [51] and ggpubr [52].

### Statistical differences between effective reproduction number of SARS-CoV-2 mutations in different levels of stringency

To determine if SARS-CoV-2 with HF_N501Y_ and not-HF_N501Y_ mutations presented statistical differences in Rt in different levels of stringency, we categorize the stringency index in ten levels: 0-10 = 0, 11-20 = 1, 21-30 = 2, 31-40 =3, 41-50 = 4, 51-60 = 5, 61-70 = 6, 71-80 = 7, 81-90 = 8, and 91-100 = 9. We estimated the distribution of the effective reproduction number 14 days after the level of stringency was implemented in each level of stringency by bootstrap using 1000 replicates. Level of stringency with at least 10 Rt points were considered in the bootstrap analysis. We also used bootstrap methods to estimate the distribution of the difference of the Rt assuming that both Rt (HF_N501Y_ and not-HF_N501Y_) comes from the same distribution and calculate the p-value of the observed difference.

### Calculation of change in time of the effective reproduction number of SARS-CoV-2 mutations

Change of Rt was calculated by subtracting the value of Rt fourteen days after the day of interest with the mean of the Rt from the day of interest to thirteen days after the day of interest.

## Author contributions

SJA designed the study. SJA, DZS, CHR, GLB, ACC, RG-SC, CSUC, RPC analyzed the data. SJA, CSUC GU-C wrote python and R scripts. The manuscript was written by SJA, DZS, CHR, GLB, ACC, RG-SC, CSUC. All authors discussed the methodologies, results, and read and approved the manuscript.

## Acknowledgements

We are very grateful to the GISAID Initiative and all its data contributors, i.e., the authors from the Originating laboratories responsible for obtaining the specimens and the Submitting laboratories where genetic sequence data were generated and shared via the GISAID Initiative, on which this research is based.

We thank Prof. Jose Luis Mena (Faculty of Biological Sciences – Universidad Ricardo Palma) and PhD(c). Carlos Prete (Escola Politecnica – University of Sao Paulo) for their valuable help in the epidemiological and statistical analysis.

We thank Prof. Shaker Chuck Farah (Institute of Chemistry – University of Sao Paulo) and PhD(c). Charlie Whittaker (Faculty of Medicine – Imperial College London) for English writing corrections and helpful comments.

To the Ricardo Palma University High-Performance Computational Cluster (URPHPC) managers Gustavo Adolfo Abarca Valdiviezo and Roxana Paola Mier Hermoza at the Ricardo Palma Informatic Department (OFICIC) for their contribution in programs and remote use configuration of URPHPC. Also, to one of the managers of the High-Performance Computer from the Instituto de Investigaciones de la Amazonía Peruana, Rodolfo Cardena Vigo for its assistance in configurations and program installations.

## Data availability

Publicly available datasets were analyzed in this study. This data can be found at: gisaid.org. All the code used to perform the analysis of this manuscript is publicly available in: https://github.com/sanjusare/Justo_et_al_2021_SR

