## Supplemental Figures and Tables for "Dynamics of SARS-CoV-2 mutations reveals regional-specificity and similar trends of N501 and High-Frequency mutation N501Y in different levels of control measures"

**Supplementary File**

Santiago Justo Arevalo*^1,2^, Daniela Zapata Sifuentes^1^, César J. Huallpa^3^, Gianfranco Landa Bianchi^1^, Adriana Castillo Chávez^1^, Romina Garavito-Salini Casas^1^, Carmen Sofia Uribe Calampa^1^, Guillermo Uceda-Campos^4^, Roberto Pineda Chavarría^1^

^1^Facultad de Ciencias Biológicas, Universidad Ricardo Palma, Lima, Peru

^2^Department of Biochemistry, Institute of Chemistry, University of São Paulo, São Paulo, Brazil

^3^Facultad de Ciencias, Universidad Nacional Agraria La Molina, Lima, Peru

^4^Facultad de Ciencias Biológicas, Universidad Nacional Pedro Ruiz Gallo, Lambayeque, Peru

**Table 1.** 115 mutations estimated to be present in more than 3 % global COVID-19 cases. ID: code of identification on the format: nucleotide change_aminoacid change_region of mutation. NRFp: Global normalized by COVID-19 cases relative frequency. Type: type of temporal dynamic of the mutation; High-Frequency (HF), Medium-Frequency (MF), Low-Frequency (LF). nt_change: change at nucleotide level. AA_change: change at aminoacid level. Location: Specific genomic region where the mutation occurred. Syn: Synonymous change

**Supplementary Figure 1.** Normalized Relative Frequency (NRFp) of the 115 mutations with more of 0.03 NRFp.

**Supplementary Figure 2.** Temporal dynamics of High-Frequency (HF) SARS-CoV-2 mutations. Normalized by cases Relative Frequency (NRFp) of the HF mutations by month. To improve visualization mutations were divided into three columns and two rows. Left, medium and right columns show non-synonymous in non-structural proteins, non-synonymous in structural proteins, and synonymous mutations, respectively. Upper and bottom rows divide mutations according their slopes to improve visualization.

**Supplementary Figure 3.** Temporal dynamics of Medium-Frequency (MF) and Low-Frequency (LF) SARS-CoV-2 mutations. Normalized by cases Relative Frequency (NRFp) of the MF (upper row) and LF (bottom row) mutations by month. To improve visualization mutations were divided into three columns. Left, medium and right columns show non-synonymous in non-structural proteins, non-synonymous in structural proteins, and synonymous mutations, respectively.

**Supplementary Figure 4.** Normalized by cases Relative Frequency (NRFp) of the HF mutations by month separated by regions (green = Africa, red = Asia, blue = Europe, grey = North America, purple = Oceania, yellow= South America). Graphics are also separated by groups and subtypes according to their similarity in the dynamics in the six regions.

**Supplementary Figure 5.** Normalized by cases Relative Frequency (NRFp) of the HF mutations by month separated by regions (green = Africa, red = Asia, blue = Europe, grey = North America, purple = Oceania, yellow= South America). Graphics are also separated by groups and subtypes according to their similarity in the dynamics in the six regions.

**Supplementary Figure 6.** Normalized by cases Relative Frequency (NRFp) of the MF mutations by month separated by regions (green = Africa, red = Asia, blue = Europe, grey = North America, purple = Oceania, yellow= South America). Graphics are also separated by subtypes according to their similarity in the dynamics in the six regions.

**Supplementary Figure 7.** Relative frequencies by month separated by countries of the MF mutations with region-specific patterns. Two first columns show mutations characteristic from South America, and the two last columns show mutations characteristic from North America.

**Supplementary Figure 8.** Relative frequencies by month separated by countries of the MF mutation with region-specificity to Europe.

**Supplementary Figure 9.** Cumulative number of cases of MF mutations with region-specificity estimated for each country with at least one case of the mutation. Two first columns show mutations characteristic from South America, and the two last columns show mutations characteristic from North America

**Supplementary Figure 10.** Cumulative number of cases of MF mutations with specificity for Europe estimated for each country with at least one case of the mutation.

**Supplementary Figure 11.** Relative frequencies by month separated by countries of the MF mutations with region-specific patterns and with a decline in the last months using data from August 10^th^ 2021. The first column shows a mutation characteristic from South America, and the last column shows a mutation characteristic from North America.

**Supplementary Figure 12.** Relative frequencies by month separated by countries of the MF mutations with region-specificity to Europe and with a decline in the last months using data from August 10^th^ 2021.

**Supplementary Figure 13.** Normalized by cases Relative Frequency (NRFp) of the LF mutations by month separated by regions (green = Africa, red = Asia, blue = Europe, grey = North America, purple = Oceania, yellow= South America). Graphics are also separated by subtypes according to their similarity in the dynamics in the six regions.

**Supplementary Figure 14.** LF mutations are region-specific or have mid-frequencies in several regions. (left-column) Normalized by cases Relative Frequency (NRFp) of the mutations by month separated by regions (green = Africa, red = Asia, blue = Europe, grey = North America, purple = Oceania, yellow= South America). (middle-column) Total NRFp by region of the analyzed Low-Frequency mutations. Numbers in each bar represents the estimated total number of cases of the particular mutation in that region. (right-column) Chi-square p-value and Pearson residuals analysis of Medium-Frequency mutations. Upper line corresponds to the mutant state and the bottom line to the not-mutant state. Grey and red boxes mean negative or positive association with the state, respectively. Intensity of the colors means higher residuals that means greater contribution.

**Supplementary Figure 15.** Effective reproduction number (Rt) of MF_R203K_ and not-MF_R203K_ are sometimes similar and sometimes different. **(a)** Each panel shows the estimated effective reproduction number of SARS-CoV-2 bearing (green) or not (grey) the MF mutation R203K (MF or notMF, respectively) in different countries. Grey bars are showing the level of stringency. Shades show a 97.5 % confidence interval in the estimation of Rt. **(b)** Statistical comparison between the bootstrap distribution of the Rt of SARS-CoV-2 bearing (blue) or not (grey) the MF mutation R203K (MF or notMF, respectively) in different levels of stringency. Points represent the mean and the lines represent the 25 and 75 percentiles of the bootstrap distribution. ** means p-value lesser than 0.05 and ns means p-value higher than 0.05.

**Table 1**

| ID | NRFp | Type | nt_change | AA_change | Location |
| --- | --- | --- | --- | --- | --- |
| G210T_---_5-UTR | 0.033 | LF | G210T | --- | 5-UTR |
| C241T_---_5-UTR | 0.971 | HF | C241T | --- | 5-UTR |
| C313T_Syn_nsp1 | 0.031 | LF | C313T | Syn | nsp1 |
| T445C_Syn_nsp1 | 0.07 | MF | T445C | Syn | nsp1 |
| T733C_Syn_nsp1 | 0.034 | LF | T733C | Syn | nsp1 |
| C913T_Syn_nsp2 | 0.136 | HF | C913T | Syn | nsp2 |
| C1059T_T85I_nsp2 | 0.194 | MF | C1059T | T85I | nsp2 |
| C2749T_Syn_nsp3 | 0.034 | LF | C2749T | Syn | nsp3 |
| C3037T_Syn_nsp3 | 0.978 | HF | C3037T | Syn | nsp3 |
| C3267T_T183I_nsp3 | 0.153 | HF | C3267T | T183I | nsp3 |
| C3828T_S370L_nsp3 | 0.035 | LF | C3828T | S370L | nsp3 |
| C4543T_Syn_nsp3 | 0.032 | MF | C4543T | Syn | nsp3 |
| C5388A_A890N_nsp3 | 0.136 | HF | C5388A | A890N | nsp3 |
| A5648C_K977Q_nsp3 | 0.034 | LF | A5648C | K977Q | nsp3 |
| C5986T_Syn_nsp3 | 0.138 | HF | C5986T | Syn | nsp3 |
| C6286T_Syn_nsp3 | 0.072 | MF | C6286T | Syn | nsp3 |
| A6319G_Syn_nsp3 | 0.037 | LF | A6319G | Syn | nsp3 |
| A6613G_Syn_nsp3 | 0.035 | LF | A6613G | Syn | nsp3 |
| T6954C_I1412T_nsp3 | 0.138 | HF | T6954C | I1412T | nsp3 |
| G8083A_M1788I_nsp3 | 0.038 | LF | G8083A | M1788I | nsp3 |
| C10319T_L89F_nsp5 | 0.097 | LF | C10319T | L89F | nsp5 |
| G11083T_L37F_nsp6 | 0.037 | MF | G11083T | L37F | nsp6 |
| T11288-_Δ106-108_nsp6 | 0.175 | HF | T11288- | Δ106-108 | nsp6 |
| C11289-_Δ106-108_nsp6 | 0.175 | HF | C11289- | Δ106-108 | nsp6 |
| T11290-_Δ106-108_nsp6 | 0.175 | HF | T11290- | Δ106-108 | nsp6 |
| G11291-_Δ106-108_nsp6 | 0.175 | HF | G11291- | Δ106-108 | nsp6 |
| G11292-_Δ106-108_nsp6 | 0.174 | HF | G11292- | Δ106-108 | nsp6 |
| T11293-_Δ106-108_nsp6 | 0.174 | HF | T11293- | Δ106-108 | nsp6 |
| T11294-_Δ106-108_nsp6 | 0.174 | HF | T11294- | Δ106-108 | nsp6 |
| T11295-_Δ106-108_nsp6 | 0.174 | HF | T11295- | Δ106-108 | nsp6 |
| T11296-_Δ106-108_nsp6 | 0.174 | HF | T11296- | Δ106-108 | nsp6 |
| C12053T_L71F_nsp7 | 0.038 | LF | C12053T | L71F | nsp7 |
| C12778T_Syn_nsp9 | 0.034 | LF | C12778T | Syn | nsp9 |
| C13860T_Syn_nsp12 | 0.036 | LF | C13860T | Syn | nsp12 |
| C14120T_P227L_nsp12 | 0.031 | LF | C14120T | P227L | nsp12 |
| C14408T_P323L_nsp12 | 0.98 | HF | C14408T | P323L | nsp12 |
| C14676T_Syn_nsp12 | 0.139 | HF | C14676T | Syn | nsp12 |
| C14805T_Syn_nsp12 | 0.043 | LF | C14805T | Syn | nsp12 |
| C15279T_Syn_nsp12 | 0.138 | HF | C15279T | Syn | nsp12 |
| G15766T_V776L_nsp12 | 0.032 | MF | G15766T | V776L | nsp12 |
| T16176C_Syn_nsp12 | 0.137 | HF | T16176C | Syn | nsp12 |
| G17259T_E341D_nsp13 | 0.036 | LF | G17259T | E341D | nsp13 |
| A17615G_K460R_nsp13 | 0.034 | MF | A17615G | K460R | nsp13 |
| A18424G_N129D_nsp14 | 0.088 | LF | A18424G | N129D | nsp14 |
| C18877T_Syn_nsp14 | 0.089 | MF | C18877T | Syn | nsp14 |
| T19839C_Syn_nsp15 | 0.036 | LF | T19839C | Syn | nsp15 |
| A20268G_Syn_nsp15 | 0.072 | MF | A20268G | Syn | nsp15 |
| G21255C_Syn_nsp16 | 0.067 | MF | G21255C | Syn | nsp16 |
| C21304T_Syn_nsp16 | 0.082 | LF | C21304T | Syn | nsp16 |
| C21614T_L18F_S | 0.059 | MF | C21614T | L18F | S |
| C21621A_T20N_S | 0.034 | LF | C21621A | T20N | S |
| C21638T_P26S_S | 0.037 | LF | C21638T | P26S | S |
| T21765-_Δ69-70_S | 0.147 | HF | T21765- | Δ69-70 | S |
| A21766-_Δ69-70_S | 0.145 | HF | A21766- | Δ69-70 | S |
| C21767-_Δ69-70_S | 0.145 | HF | C21767- | Δ69-70 | S |
| A21768-_Δ69-70_S | 0.145 | HF | A21768- | Δ69-70 | S |
| T21769-_Δ69-70_S | 0.145 | HF | T21769- | Δ69-70 | S |
| G21770-_Δ69-70_S | 0.145 | HF | G21770- | Δ69-70 | S |
| G21974T_D138Y_S | 0.046 | LF | G21974T | D138Y | S |
| T21991-_Δ144_S | 0.134 | HF | T21991- | Δ144 | S |
| T21992-_Δ144_S | 0.134 | HF | T21992- | Δ144 | S |
| A21993-_Δ144_S | 0.135 | HF | A21993- | Δ144 | S |
| G22132T_R190S_S | 0.033 | LF | G22132T | R190S | S |
| C22227T_A222V_S | 0.074 | MF | C22227T | A222V | S |
| C22444T_D294D_S | 0.037 | LF | C22444T | D294D | S |
| T22917G_L452R_S | 0.054 | LF | T22917G | L452R | S |
| G22992A_S477N_S | 0.037 | MF | G22992A | S477N | S |
| G23012A_E484K_S | 0.08 | MF | G23012A | E484K | S |
| A23063T_N501Y_S | 0.181 | HF | A23063T | N501Y | S |
| C23271A_A570D_S | 0.136 | HF | C23271A | A570D | S |
| A23403G_D614G_S | 0.989 | HF | A23403G | D614G | S |
| C23525T_H655Y_S | 0.037 | LF | C23525T | H655Y | S |
| C23604A_P681H_S | 0.167 | HF | C23604A | P681H | S |
| C23604G_P681R_S | 0.034 | LF | C23604G | P681R | S |
| C23709T_T716I_S | 0.139 | HF | C23709T | T716I | S |
| T24506G_S982A_S | 0.137 | HF | T24506G | S982A | S |
| C24642T_T1027I_S | 0.035 | LF | C24642T | T1027I | S |
| G24914C_D1118H_S | 0.137 | HF | G24914C | D1118H | S |
| G25088T_V1176F_S | 0.082 | LF | G25088T | V1176F | S |
| G25563T_Q57H_ORF3a | 0.285 | MF | G25563T | Q57H | ORF3a |
| C25710T_L106L_ORF3a | 0.031 | MF | C25710T | L106L | ORF3a |
| G25907T_G172V_ORF3a | 0.087 | LF | G25907T | G172V | ORF3a |
| T26149C_S253P_ORF3a | 0.037 | LF | T26149C | S253P | ORF3a |
| C26681T_F53F_M | 0.047 | LF | C26681T | F53F | M |
| C26735T_Y71Y_M | 0.08 | MF | C26735T | Y71Y | M |
| C26801G_L93L_M | 0.068 | MF | C26801G | L93L | M |
| T27299C_I33T_ORF6 | 0.04 | LF | T27299C | I33T | ORF6 |
| C27944T_H17H_ORF8 | 0.035 | LF | C27944T | H17H | ORF8 |
| C27964T_S24L_ORF8 | 0.103 | LF | C27964T | S24L | ORF8 |
| C27972T_Q27stop_ORF8 | 0.139 | HF | C27972T | Q27stop | ORF8 |
| G28048T_R52I_ORF8 | 0.136 | HF | G28048T | R52I | ORF8 |
| A28095T_K68stop_ORF8 | 0.05 | MF | A28095T | K68stop | ORF8 |
| A28111G_Y73C_ORF8 | 0.138 | HF | A28111G | Y73C | ORF8 |
| G28167A_E92K_ORF8 | 0.034 | LF | G28167A | E92K | ORF8 |
| C28253T_F120F_ORF8 | 0.043 | MF | C28253T | F120F | ORF8 |
| A28271-_---_Intergenic | 0.143 | HF | A28271- | --- | Intergenic |
| G28280C_D3H_N | 0.129 | HF | G28280C | D3H | N |
| A28281T_D3V_N | 0.128 | HF | A28281T | D3V | N |
| T28282A_D3E_N | 0.129 | HF | T28282A | D3E | N |
| C28472T_P67S_N | 0.09 | LF | C28472T | P67S | N |
| C28512G_P80R_N | 0.034 | LF | C28512G | P80R | N |
| C28854T_S194L_N | 0.098 | MF | C28854T | S194L | N |
| C28869T_P199L_N | 0.102 | LF | C28869T | P199L | N |
| A28877T_S202C_N | 0.037 | LF | A28877T | S202C | N |
| G28878C_S202T_N | 0.036 | LF | G28878C | S202T | N |
| G28881A_R203K_N | 0.456 | MF | G28881A | R203K | N |
| G28882A_R203R_N | 0.455 | MF | G28882A | R203R | N |
| G28883C_G204R_N | 0.457 | MF | G28883C | G204R | N |
| C28887T_T205I_N | 0.055 | MF | C28887T | T205I | N |
| C28932T_A220V_N | 0.07 | MF | C28932T | A220V | N |
| G28975T_M234I_N | 0.046 | LF | G28975T | M234I | N |
| C28977T_S235F_N | 0.141 | HF | C28977T | S235F | N |
| T29148C_I292T_N | 0.036 | LF | T29148C | I292T | N |
| G29402T_D377Y_N | 0.052 | LF | G29402T | D377Y | N |
| G29645T_V30L_ORF10 | 0.07 | MF | G29645T | V30L | ORF10 |

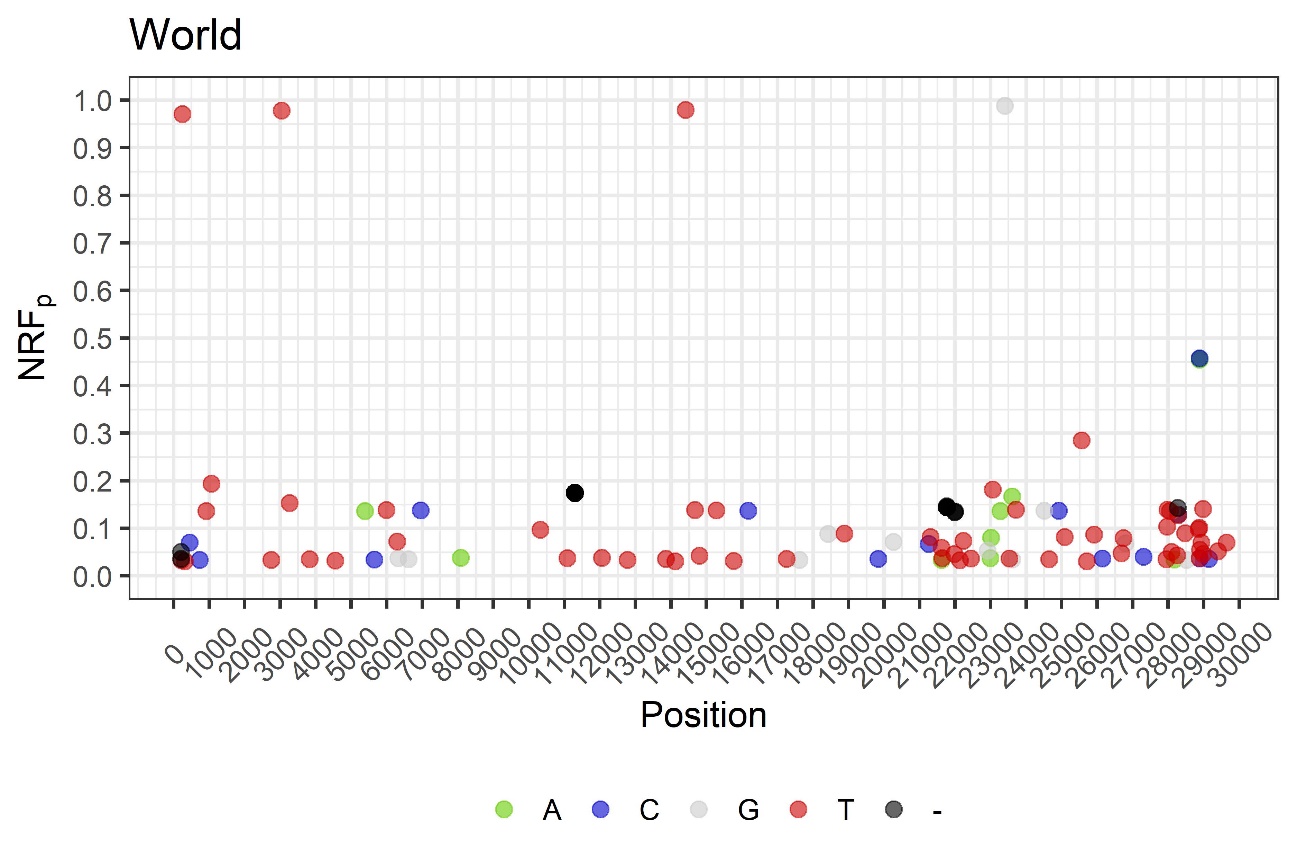

**S1**

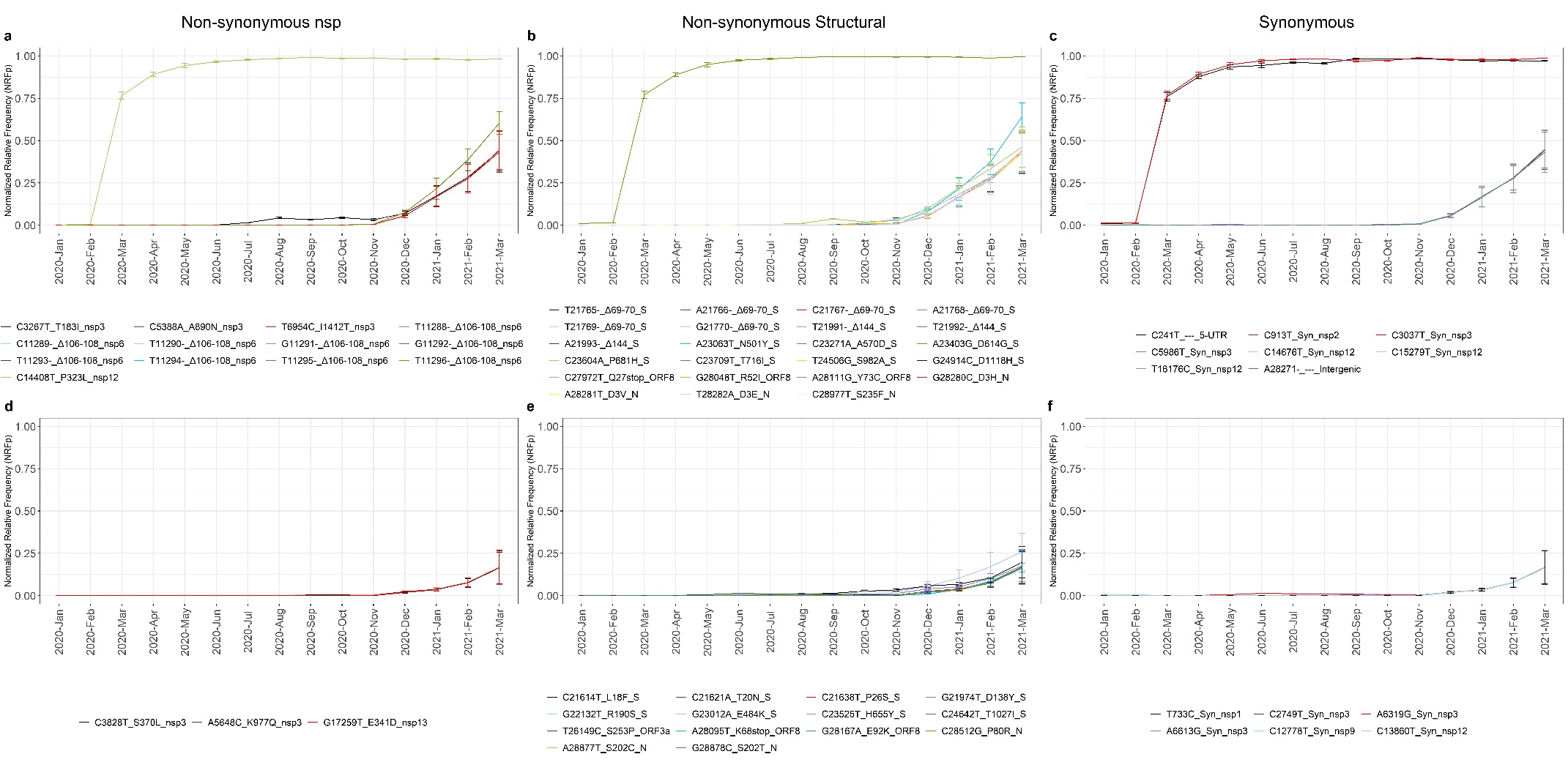

S2

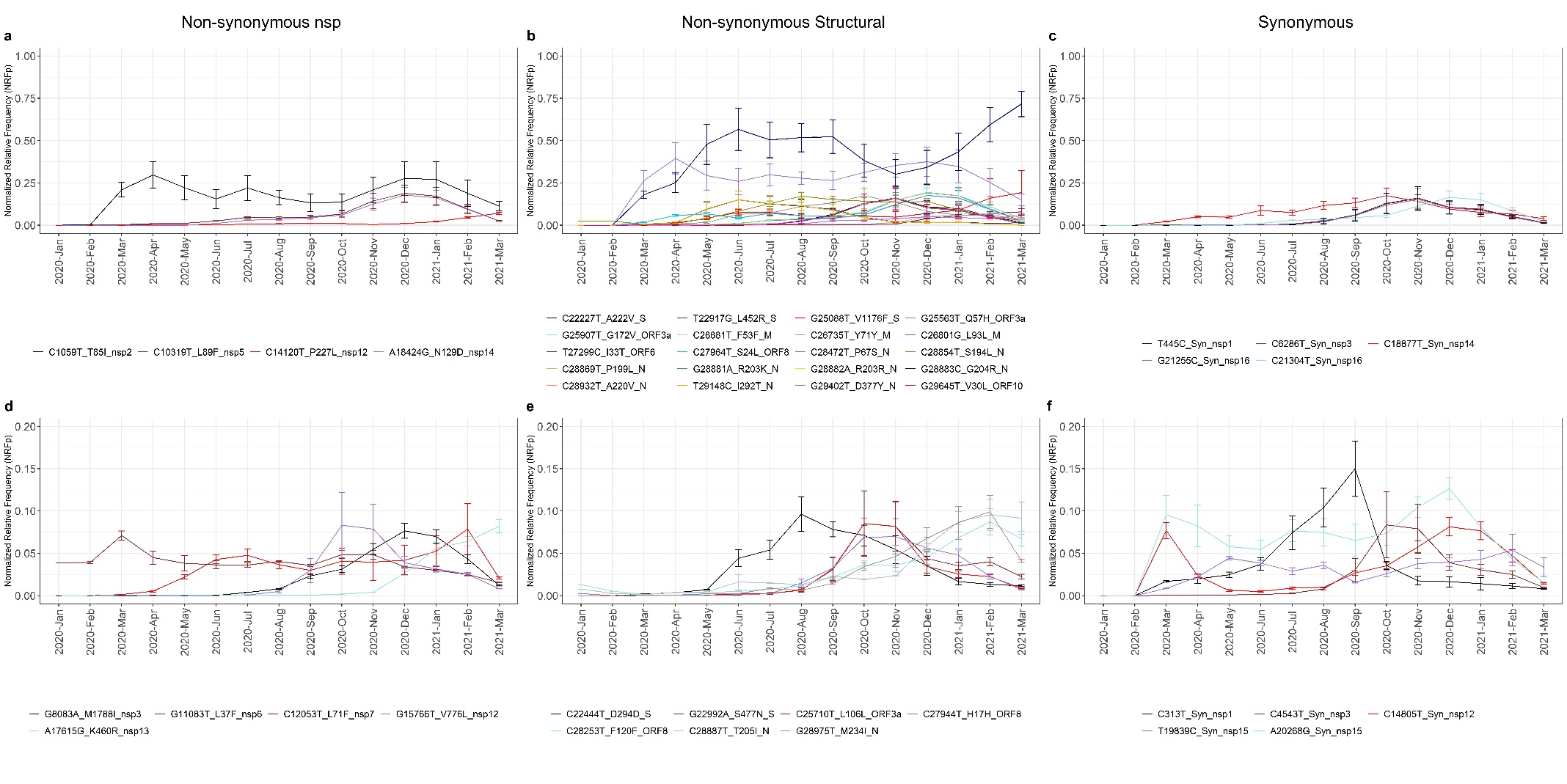

S3

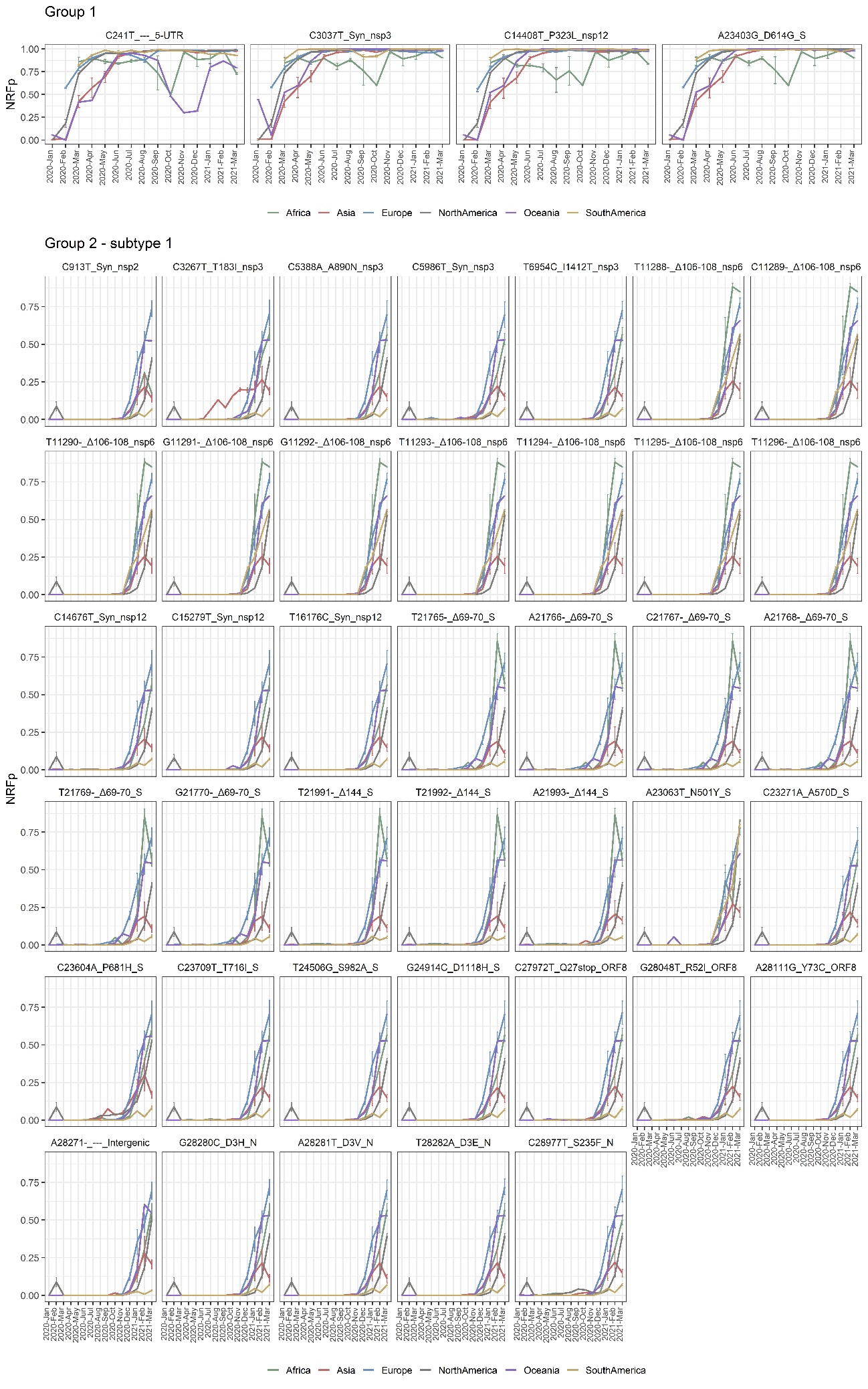

S4

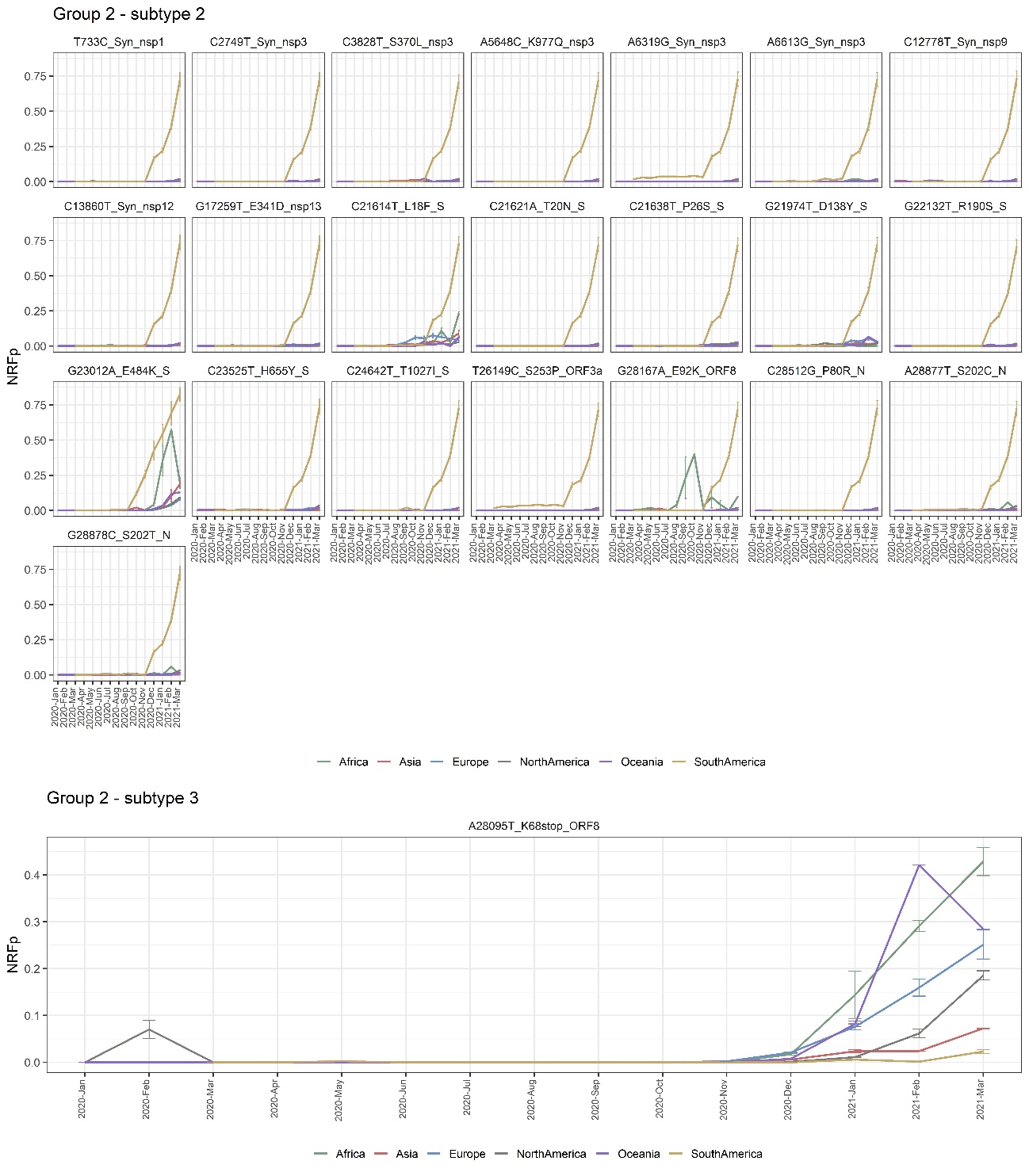

S5

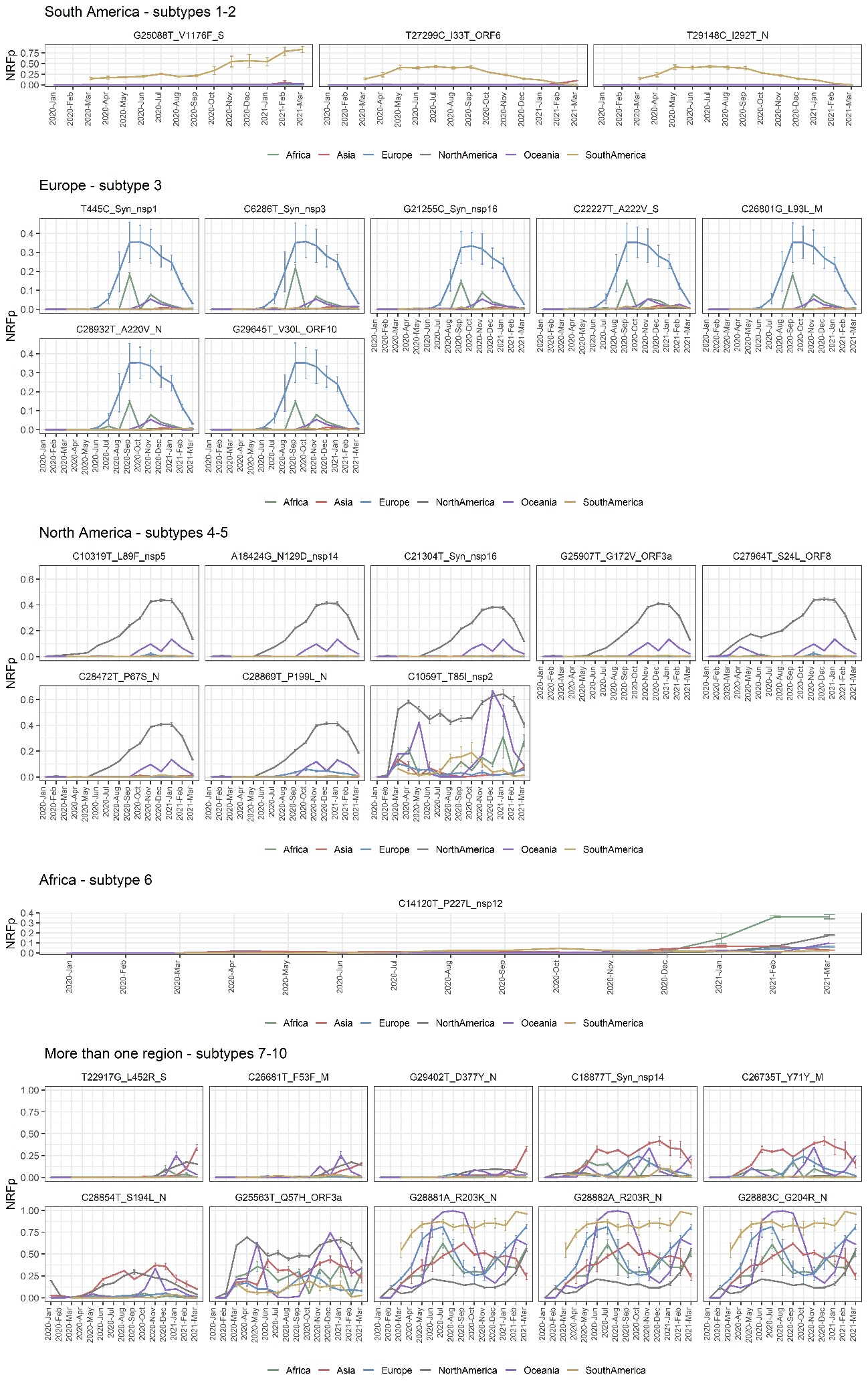

S6

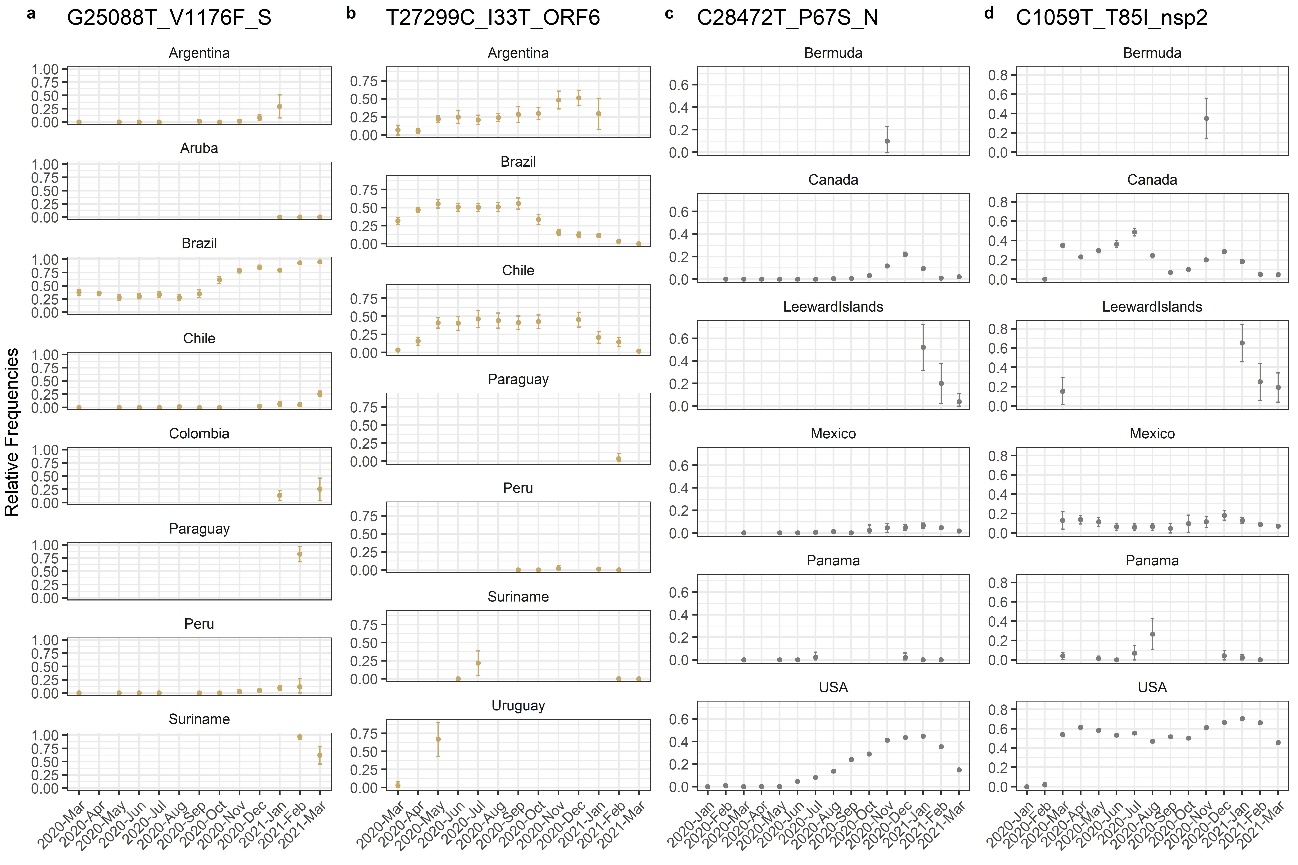

S7

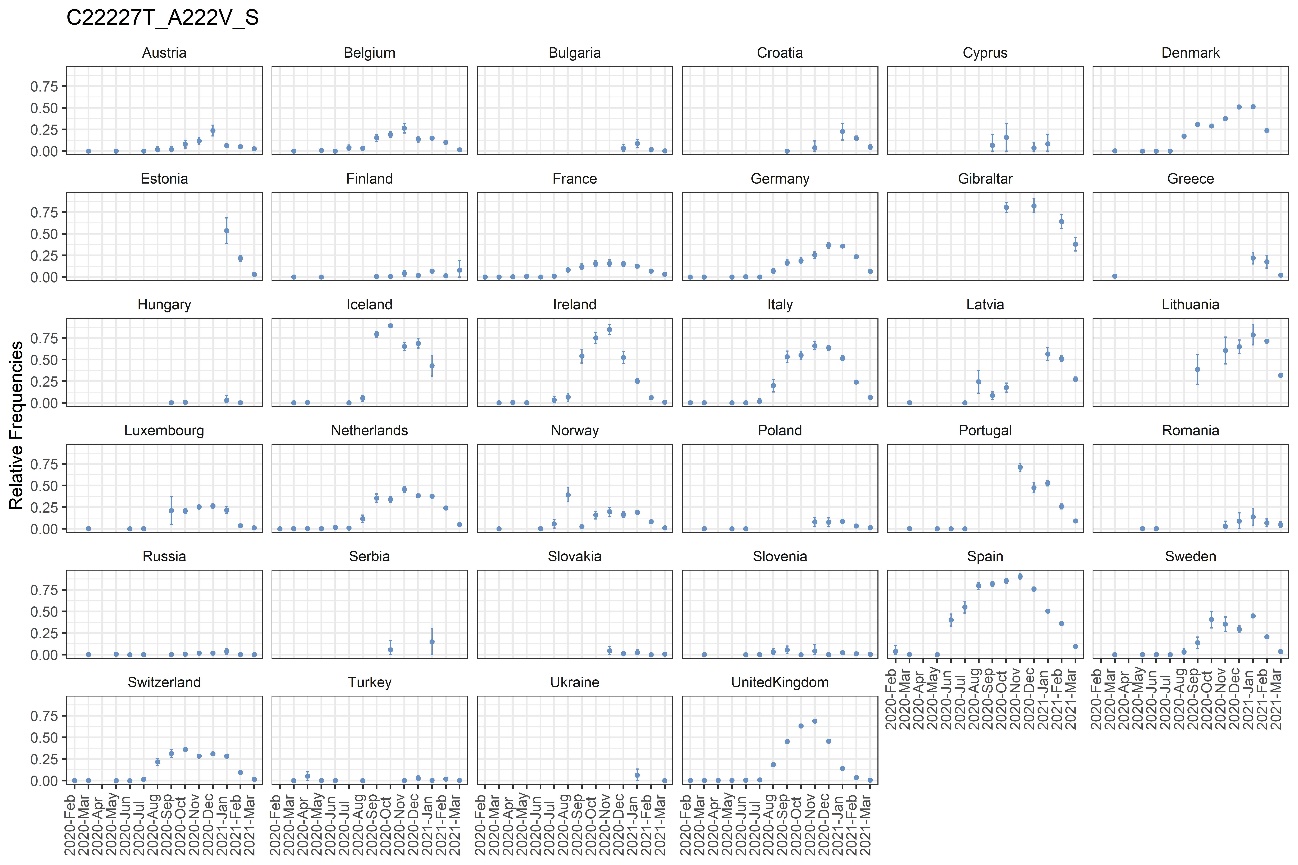

S8

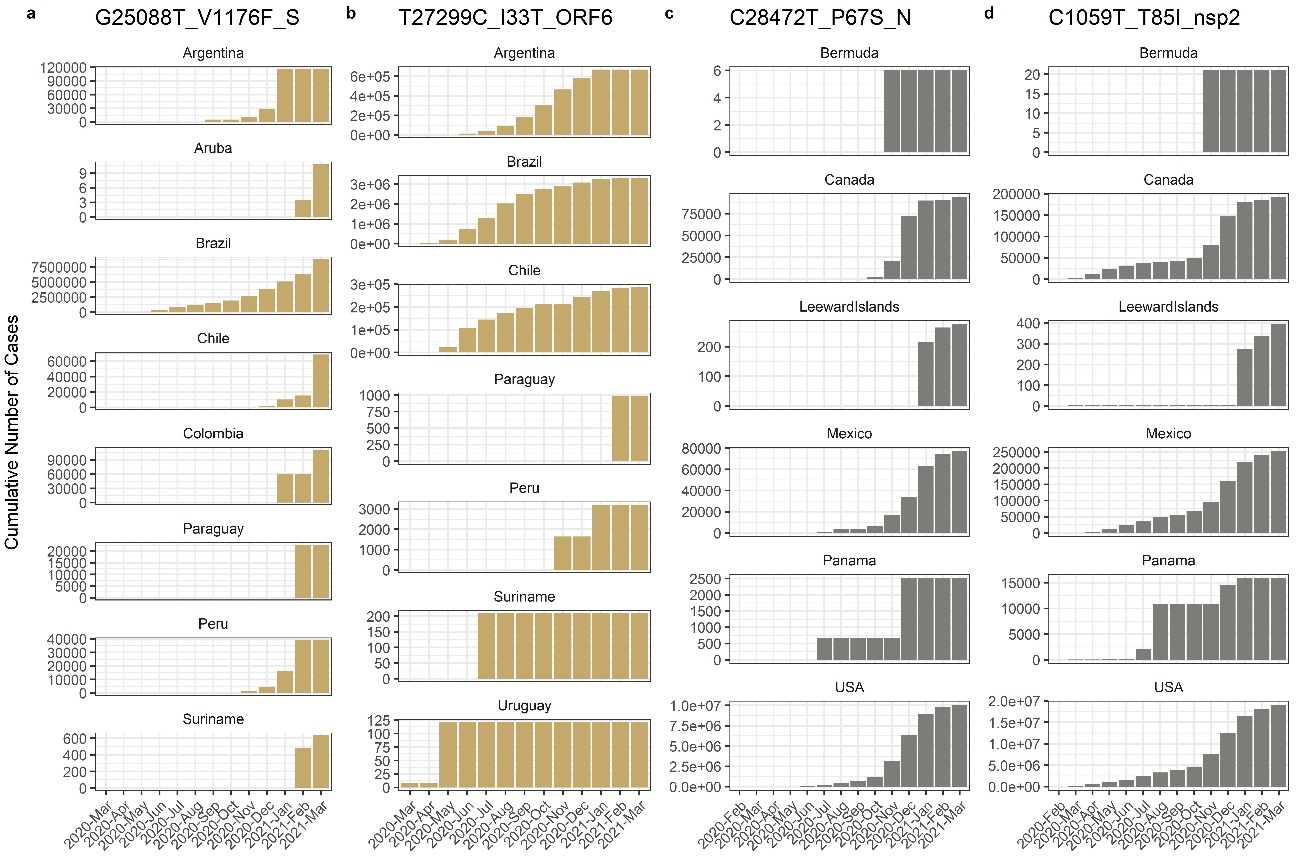

S9

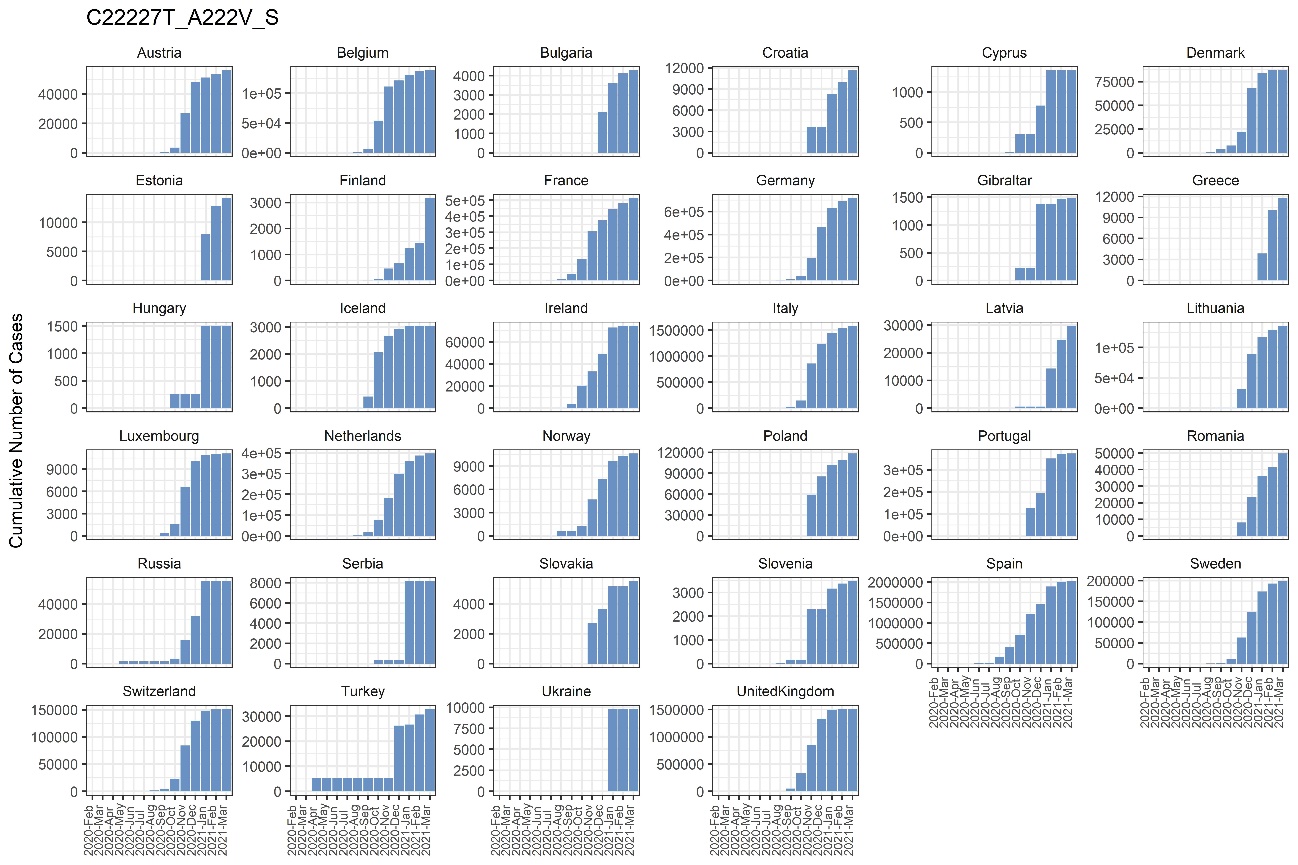

S10

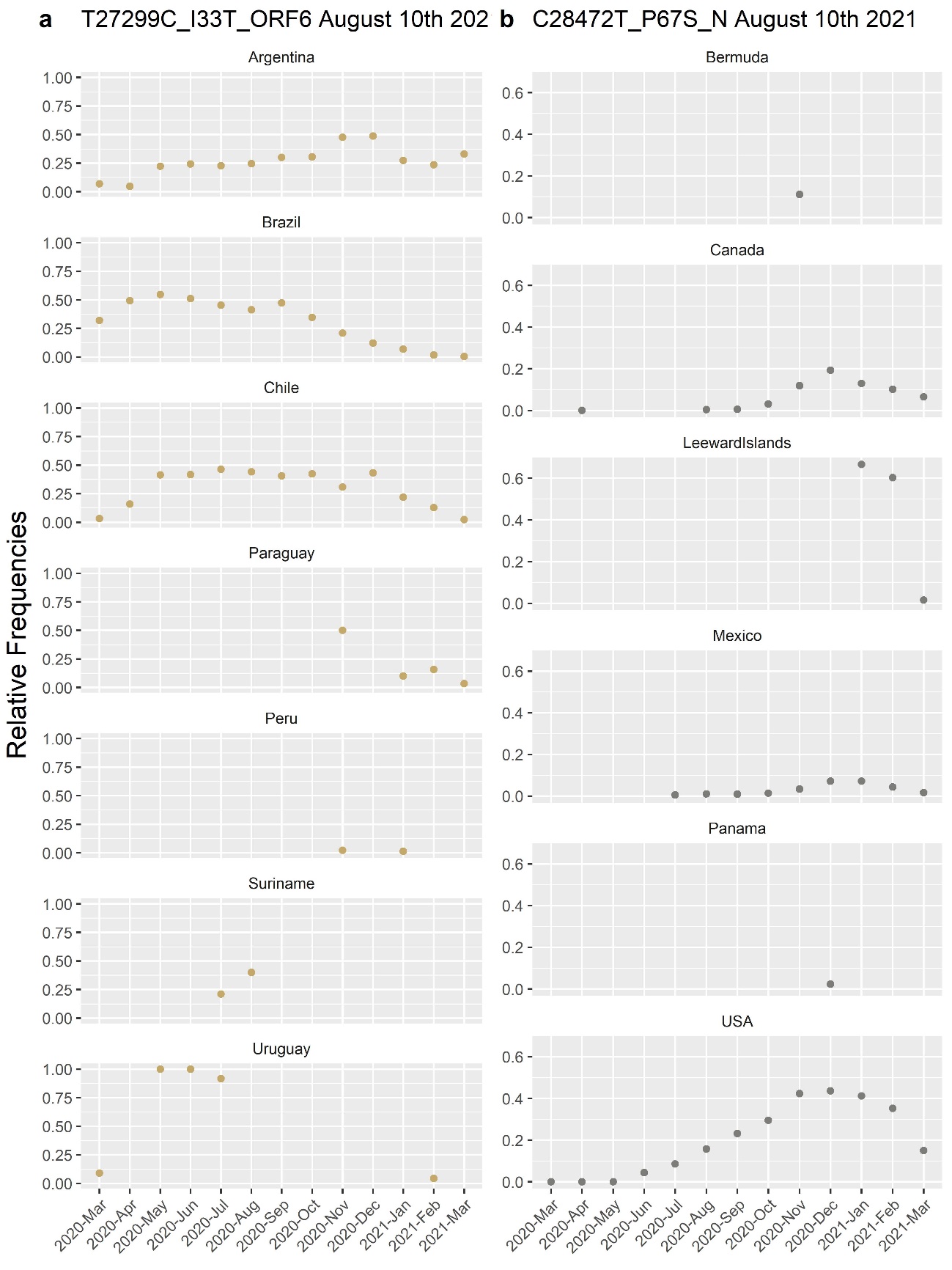

S11

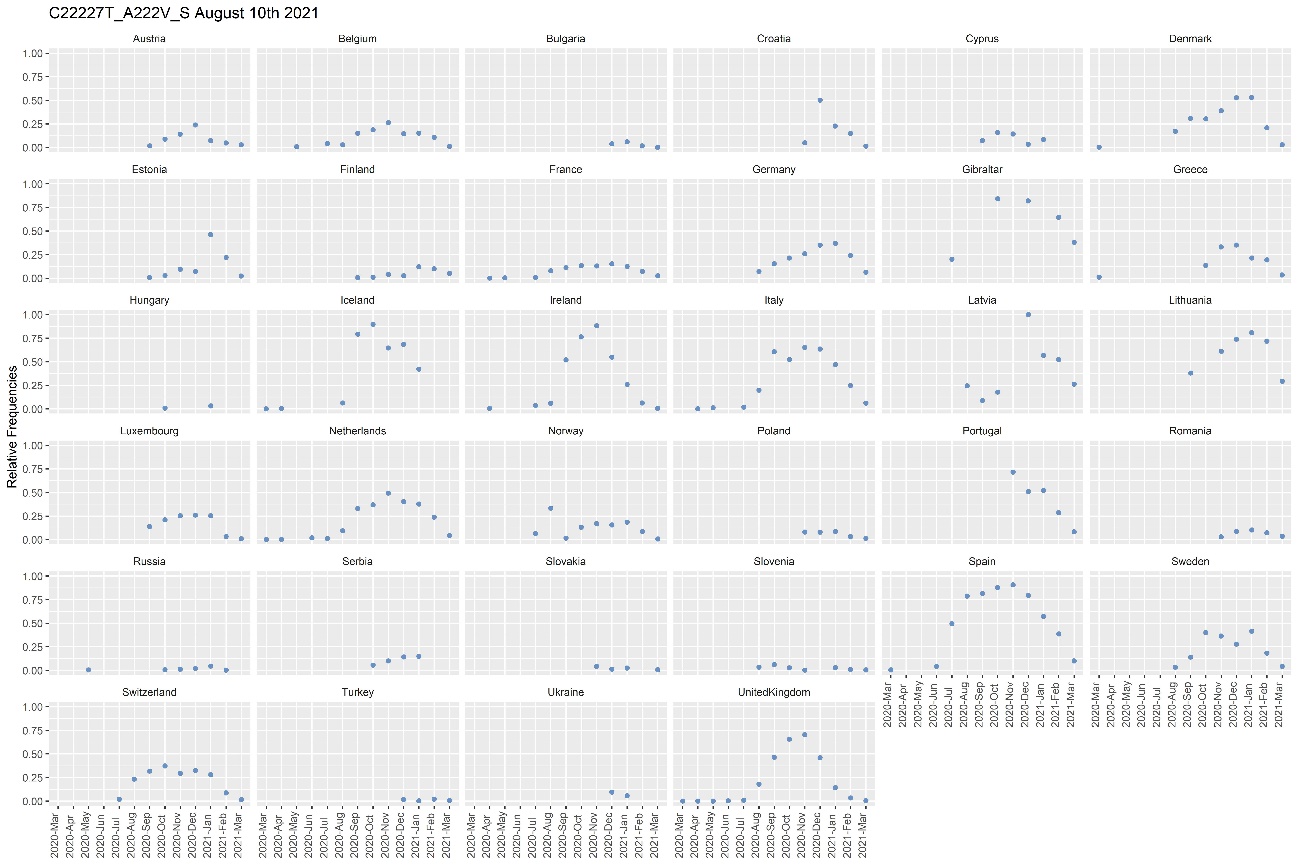

S12

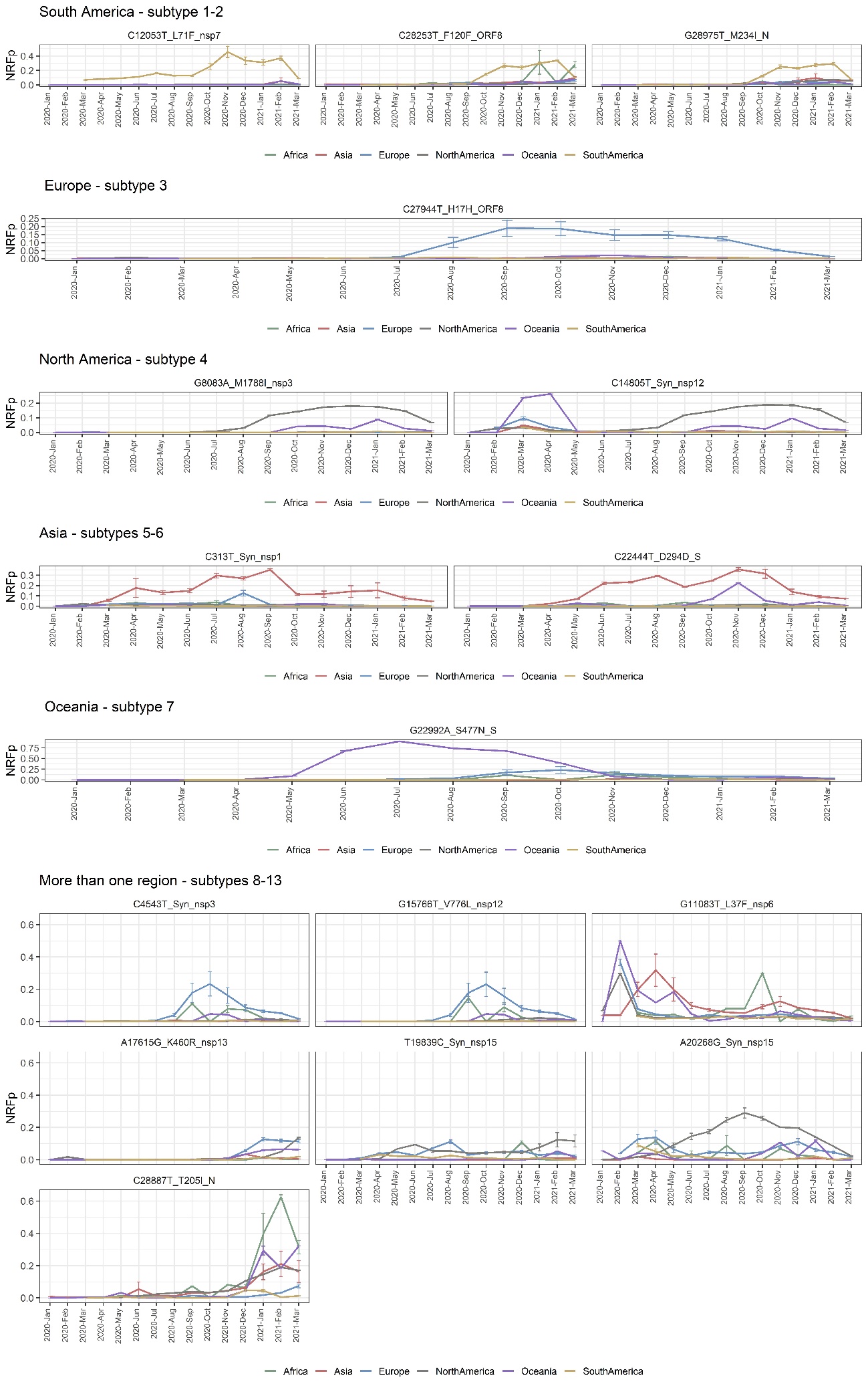

S13

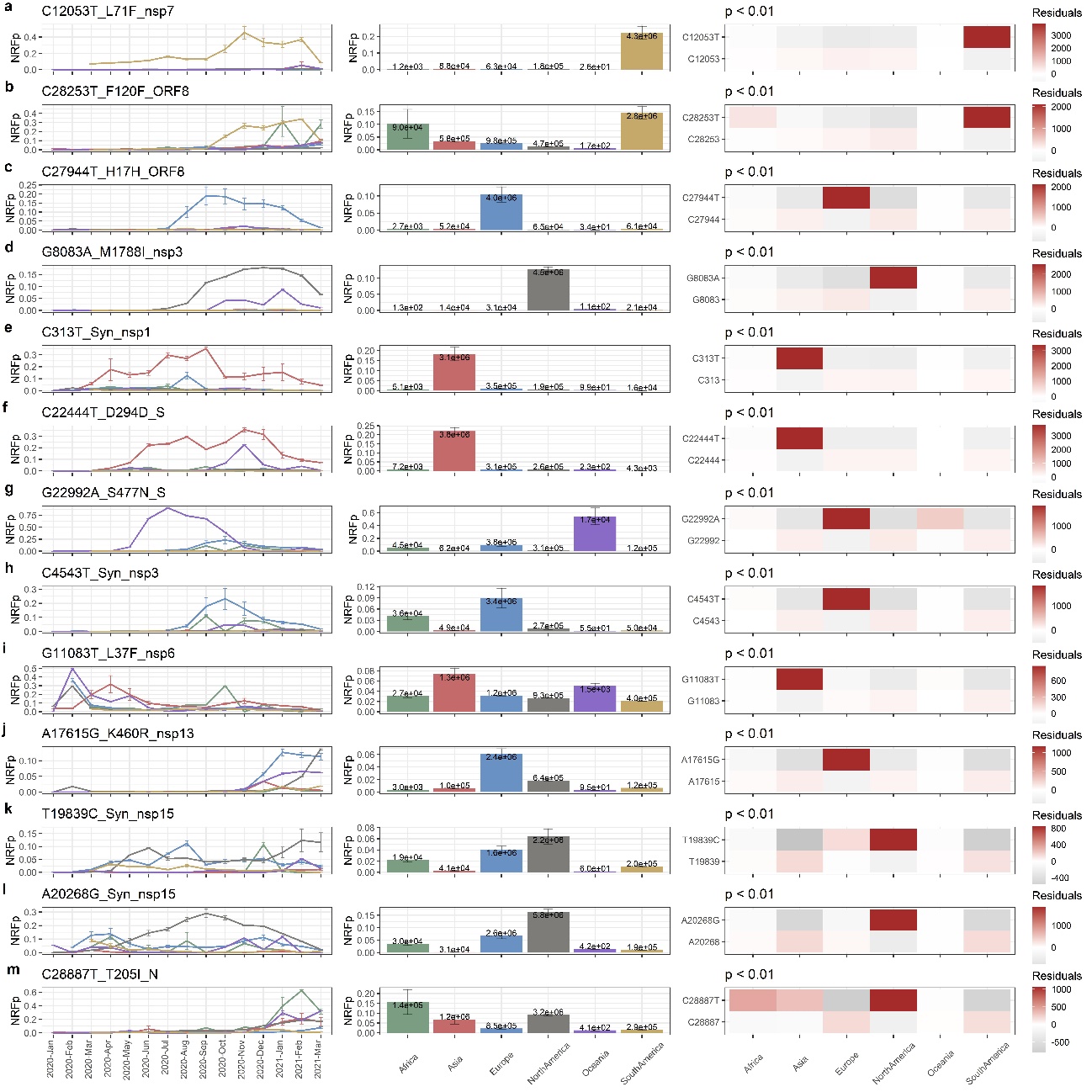

S14

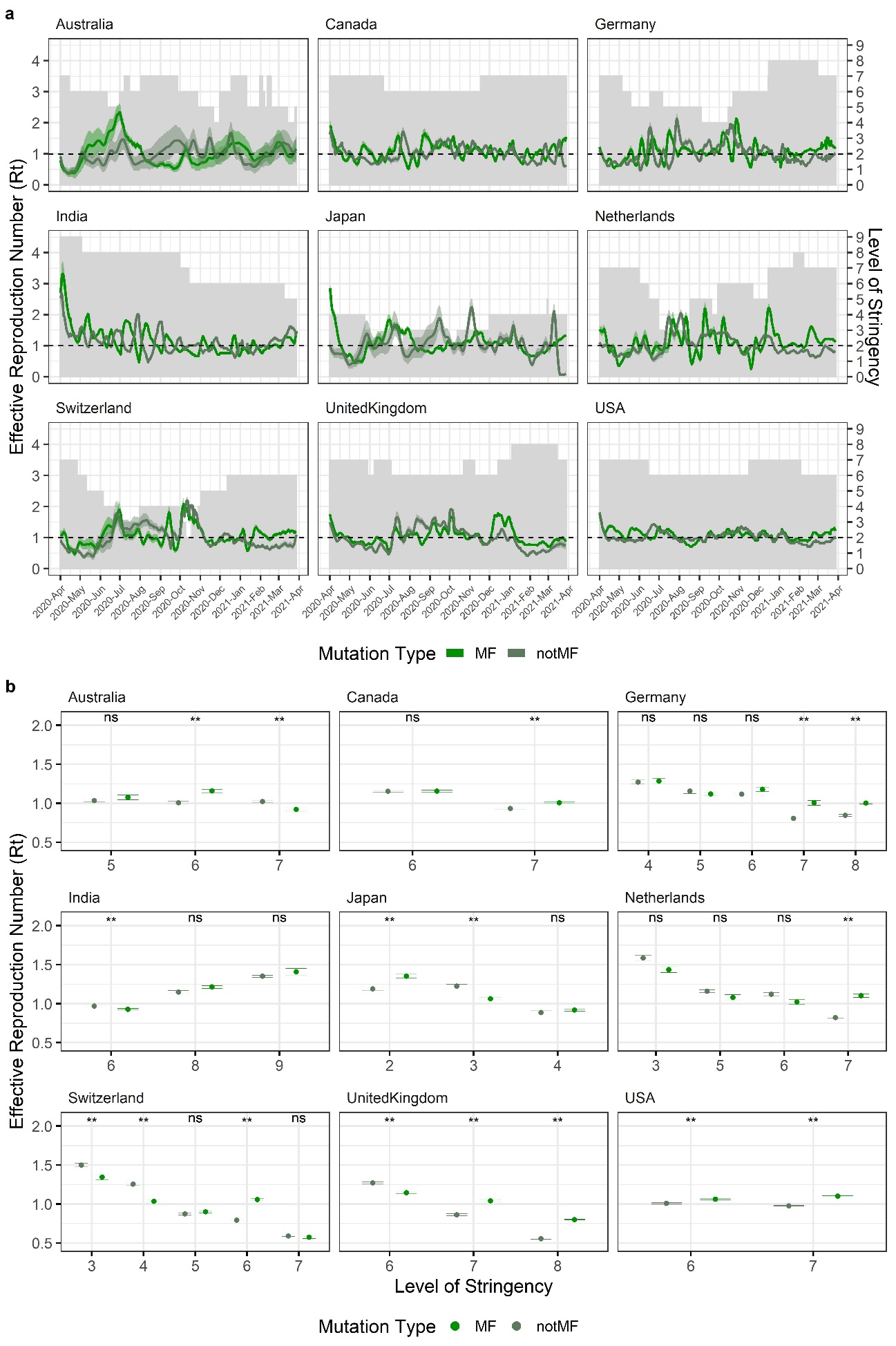

S15
